# *In vitro* engineering of the lung alveolus

**DOI:** 10.1101/2022.03.13.484143

**Authors:** Katherine L. Leiby, Yifan Yuan, Ronald Ng, Micha Sam Brickman Raredon, Taylor S. Adams, Pavlina Baevova, Karen K. Hirschi, Stuart G. Campbell, Naftali Kaminski, Erica L. Herzog, Laura E. Niklason

## Abstract

Therapeutic lung regeneration is predicated upon successful reconstitution of lung alveoli, the functional units of gas exchange. Here, we identify requisite multimodal cues that are critical to reconstructing the alveolar epithelium and alveoli in lung scaffolds. Alveolar reconstruction *in vitro* is divided into several distinct phases. In the first phase, endothelial cells coordinate with fibroblasts and select exogenous factors to promote alveolar scaffold population with surfactant-secreting alveolar epithelial type 2 cells (AEC2s). After formation of organized epithelial alveolar-like structures, subsequent withdrawal of Wnt and FGF agonism synergizes with tidal-level mechanical strain to induce differentiation of AEC2s to squamous type 1 AECs (AEC1s) in cultured alveoli, *in situ*. These results outline a rational strategy to engineer an alveolus of AEC2s and AEC1s contained within epithelial-mesenchymal-endothelial units, and reveal the critical interplay amongst biochemical, cellular, and mechanical niche cues within the reconstituting alveolus.

---

End-stage lung disease remains a leading cause of death in the United States^1^. The only definitive therapy is lung transplantation; however, suitable donor organs are scarce and graft survival at 10 years is approximately 30%^2^. Lung acellular extracellular matrix (ECM) scaffolds provide a pre-made, bioactive template for cellular population and regeneration of functional alveolar networks, which may one day support therapeutic transplantation^3, 4^. But despite some reported progress in bioengineered lung revascularization^5, 6^, organized epithelial recellularization to reconstitute alveoli with native-like phenotypic features has been an elusive goal. The alveolar epithelium forms a gas-diffusible surface, provides the tightest barrier to fluid leak, and secretes surfactant to prevent alveolar collapse^7^, and as such is indispensable to regenerating functional lung tissue capable of gas exchange.

Alveolar epithelial type 2 cells (AEC2s) are the principal epithelial stem cell of the distal lung^8–10^. Following alveolar injury, AEC2s proliferate, and then differentiate to type 1 AECs (AEC1s) to reinstate epithelial integrity^11^. Dysregulation of AEC2 progenitor functions precludes meaningful restoration of alveolar structure and function, as is seen in multiple lung diseases^12, 13^. Although an area of intense investigation, the signals governing AEC2 self-renewal and differentiation remain poorly understood (beyond mesenchymal-derived cues), and hence harnessing AEC2s to promote alveolar regeneration has met limited success^14, 15^. Achieving AEC1 differentiation *ex vivo* from AEC2s has been particularly challenging: common approaches such as extended time in culture^8^ or growth on glass in the presence of serum^9^ are poorly defined and ill-suited for translation.

Here, we describe a strategy to engineer multilineage lung alveolar tissue by directing the AEC2 regeneration process *ex vivo* and *in situ*, within decellularized lung scaffolds (Fig. 1A). Because AEC2s modulate their behavior via integration of complex cellular, biochemical, and physical stimuli within their alveolar niche^16^, we hypothesized that guiding AEC2 proliferation and then differentiation *in vitro* – and reconstructing the surrounding alveolus – would be facilitated by simultaneous modulation of these varied multimodal cues.

**Fig. 1:**
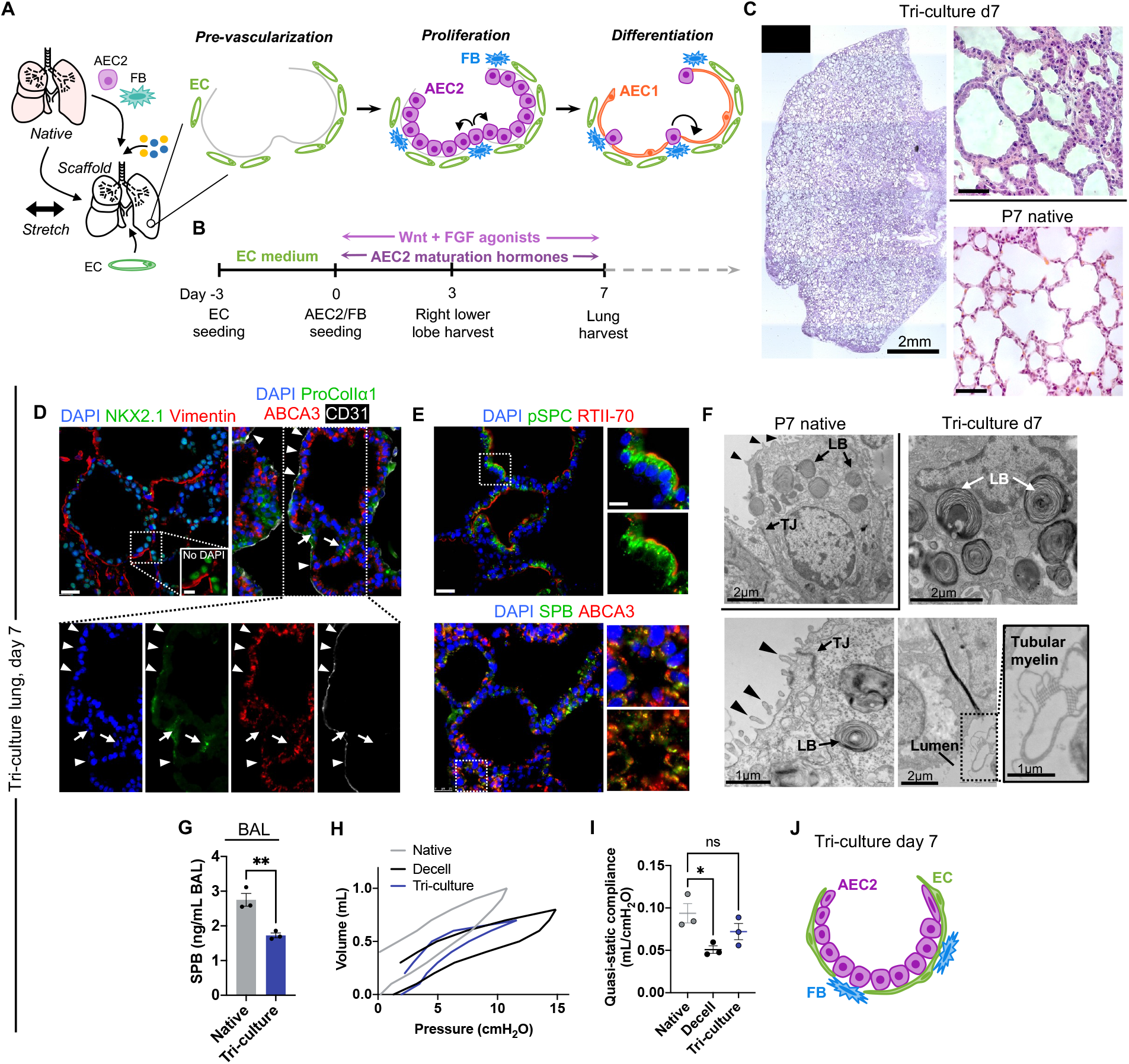
Reconstitution of a fibroblast-endothelial niche for AEC2 scaffold repopulation. (**A**) Schema for alveolar engineering. (**B**) Timeline for tri-culture lung recellularization, corresponding to pre-vascularization and the proliferation phase in (A). (**C**) H&E staining of tri-culture engineered and P7 native lungs. Scale bars, 50 μm (right). (**D**) Immunostaining of NKX2.1^+^ epithelial cells and vimentin^+^ FBs or ECs (left); or ABCA3^+^ AEC2s, procoll1α1^+^ FBs, and CD31^+^ ECs (right) in tri-culture lung. Arrowheads, CD31^+^ ECs. Arrows, procoll1α1^+^ FBs. (**E**) Immunostaining for canonical AEC2 proteins in tri-culture lung. (D and E) Scale bar, 25 μm. Magnified region, 10 μm. (**F**) TEM examining epithelial ultrastructural features in tri-culture lung, with a P7 native AEC2 shown for comparison. LB, lamellar body. TJ, tight junction. Arrowheads, apical microvilli. (**G**) Quantification of secreted SPB protein in bronchoalveolar lavage (BAL) of adult rat and tri-culture lungs (*n* = 3). Unpaired two-tailed *t*-test. (**H** and **I**) Representative quasi-static pressure-volume curves (H) with calculated compliance (I) (*n* = 3). One-way ANOVA with Holm-Sidak. (**J**) Schematic of alveolar-like structures in tri-culture lung. Error bars: mean ± SEM. ns, not significant, \**P* < 0.05, \*\**P* < 0.01.

## RESULTS

### Reconstituting a niche for AEC2 regeneration

Previous work has shown that AEC2s, when cultured alone in decellularized lung matrices, rapidly lose their differentiated epithelial phenotype^17^. We hypothesized that reintroducing critical alveolar niche constituents, beyond the lung matrix scaffold, would support AEC2 growth and phenotypic maintenance for an initial phase of alveolar repopulation. Alveolar fibroblasts (FBs), or a subset thereof, are considered the principal niche cell supporting AEC2 growth and self-renewal^11, 18^. Furthermore, homeostatic AEC2 renewal preferentially occurs in perivascular regions of the native lung^9, 19^, and endothelial cells (ECs) are required for effective engraftment of regenerative epithelium in the alveoli post-injury^20, 21^. To initially test both mesenchymal (*i.e.* fibroblastic) and endothelial support of AEC2 self-renewal, we adapted a 3-dimensional (3D) alveolosphere assay using a hydrogel substrate^8^. For alveolosphere culture, we utilized a serum-free medium containing factors supporting AEC2 growth (GSK3β inhibitor CHIR99021 [CHIR] and keratinocyte growth factor [KGF]; “CK”) and maturation (dexamethasone, cyclic AMP, 3-isobutyl-1-methylxanthine [IBMX], and retinoic acid [RA]; “DCIR”)^22, 23^. In the alveolosphere system and exposed to CK+DCIR, freshly isolated neonatal rat AEC2s (Extended Data Fig. 1) demonstrated robust alveolosphere formation when cultured together with both neonatal lung FBs (cultured for 3 days prior to seeding; Extended Data Fig. 2) and microvascular ECs (purchased from VEC Technologies) in “tri-culture” (Extended Data Fig. 3A-C). Importantly, removing either ECs or FBs from the tri-culture was associated with a significant reduction in alveolosphere numbers (Extended Data Fig. 3B,C). However, the defect associated with EC removal could be rescued by EC conditioned medium, suggesting a role for endothelial-derived soluble factors in enhancing AEC2 growth in the presence of FBs. However, FB cell contact appeared to be critical for their effects on AEC2s in the alveolosphere system (Extended Data Fig. 3B,C).

**Fig. 2.**
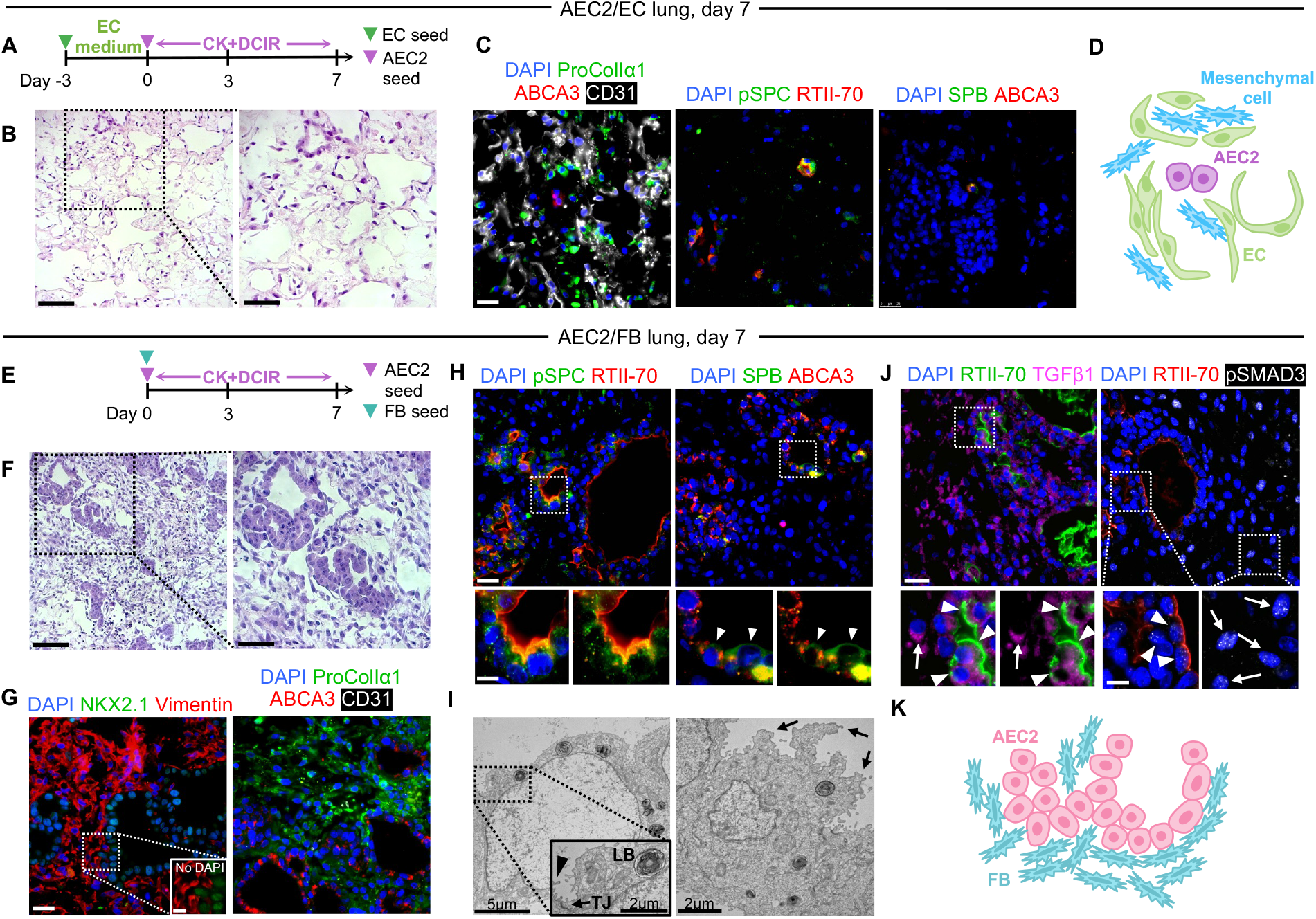
Endothelial cells and fibroblasts are insufficient as independent niche cells. (**A**) Timeline for AEC2/EC lung culture. (**B**) H&E staining of AEC2/EC lung. (**C**) Immunostaining showing few AEC2s in AEC2/EC lungs at d7. (**D**) Schematic of tissue repopulation in AEC2/EC lungs. (**E**) Timeline for AEC2/FB lung culture. (**F**) H&E staining of AEC2/FB lung. (**G**) Immunostaining of NKX2.1^+^ epithelial cells and vimentin^+^ FBs (left); or ABCA3^+^ AEC2s and procoll1α1^+^ FBs (right) in AEC2/FB lung. (**H**) Immunostaining for canonical AEC2 proteins in AEC2/FB lung. (**I**) TEM examining epithelial ultrastructure in AEC2/FB lung. LB, lamellar body. TJ, tight junction. Arrowhead, apical microvilli. Arrows, irregular apical projections. (**J**) Immunostaining showing epithelial (arrowheads) and mesenchymal (arrows) TGFβ1 and pSMAD3 expression in AEC2/FB lungs. (**K**) Schematic of alveolar-like structures in AEC2/FB lungs. Scale bars: (B and F) 100 μm. Magnified region, 50 μm. (C, G, H, and J): 25 μm. Magnified region, 10 μm.

**Fig. 3.**
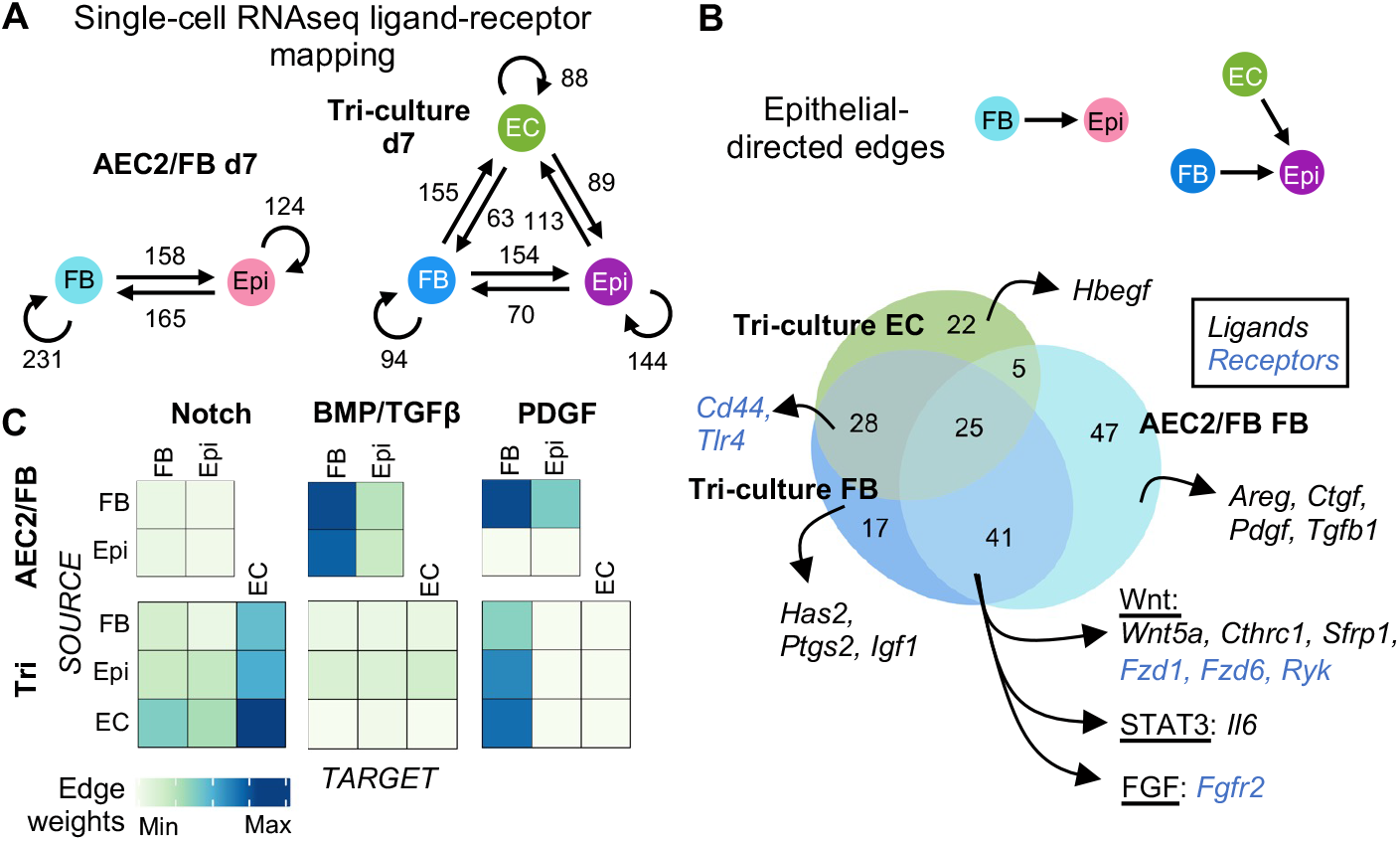
Endothelial cells alter fibroblast-AEC2 communication. (**A**) Schematic of signaling networks in AEC2/FB and tri-culture engineered lungs, from scRNAseq data. Numbers indicate the putative ligand-receptor (LR) pairs expressed along each cell-cell vector. (**B**) Venn diagram showing the overlap in ligands and receptors for FB-epi and EC-epi signaling vectors in engineered lungs. (**C**) Heatmaps of summed edge weights for all LR pairs along the given cell-cell vectors within the indicated signaling pathways in engineered lungs. Color scale applies to each pathway individually.

We next determined whether these observations applied in the more organ-mimetic system of the adult lung ECM scaffold. Acellular ECM scaffolds^24^ were seeded with ECs into the vasculature on day -3 of culture^25^, followed by seeding with AEC2s and FBs via the airway tree on day 0. Seeded scaffolds were cultured in a bioreactor^26^ with CK+DCIR medium under pulsatile vascular perfusion (Fig. 1B). The engrafted AEC2 population expanded significantly in the alveolar regions of the scaffold, and, by day 7, tri-cultured lungs were densely repopulated with a meshwork of alveolar-like structures lined by cuboidal ABCA3^+^ AEC2s (Fig. 1C,D, Extended Data Fig. 4A-D). Engineered tissue architecture was reminiscent of that of neonatal lung (Fig. 1C). The epithelial monolayers were surrounded by a thin stroma of CD31^+^ ECs and sparser procollagen Iα1^+^ FBs near the septal junctions (Fig. 1D), similar to alveolar FB localization in the native lung^27^. In contrast, tri-culture lungs cultured without CK+DCIR medium supplementation supported only rare AEC2s after 7 days, and exhibited substantial disruption of alveolar architecture (Extended Data Fig. 5).

**Fig. 4.**
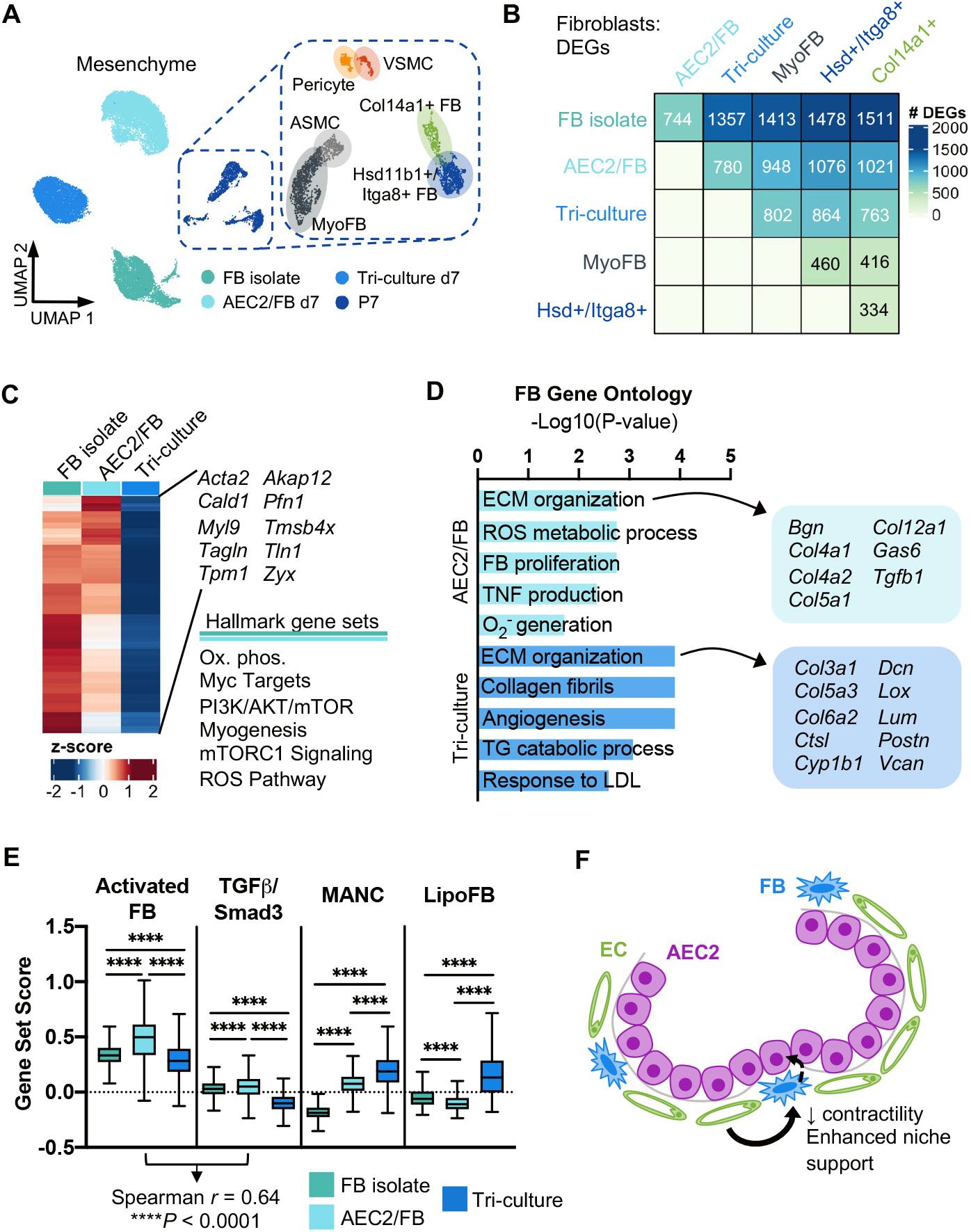
Endothelial cells enhance mesenchymal niche support. (**A**) Merged UMAP embedding of scRNAseq data for the FB isolate, engineered lung FBs, and native P7 mesenchyme. ASMC, airway smooth muscle cell. MyoFB, myofibroblast. VSMC, vascular smooth muscle cell. (**B**) Heatmap showing numbers of differentially expressed genes (DEGs) between FB subsets. (**C**) Heatmap of genes enriched in both the FB isolate and AEC2/FB FBs, as compared to tri-culture FBs, with associated pathway enrichment. (**D**) Curated biological processes enriched in AEC2/FB and tri-culture FBs. DEGs for analysis were defined as those expressed in a minimum of 25% of cells with fold change ≥2 and adjusted *P* < 0.05. DEGs associated with “ECM organization” are indicated at right. (**E**) Scoring of individual FBs for the indicated signatures. Scores > 0 indicate enriched expression compared to random gene sets. Correlation *r* between activated FB and TGFβ/Smad3 scores in co-culture FBs is indicated. MANC, mesenchymal alveolar niche cell. LipoFB, lipofibroblast. Boxplots: center indicates median, box limits indicate 1^st^ and 3^rd^ quartiles, whiskers indicate min and max (outliers not shown). Kruskal-Wallis test with Dunn’s post-test. \*\*\*\**P* < 0.0001. (**F**) Proposed model for endothelial indirect support of AEC2s.

**Fig. 5.**
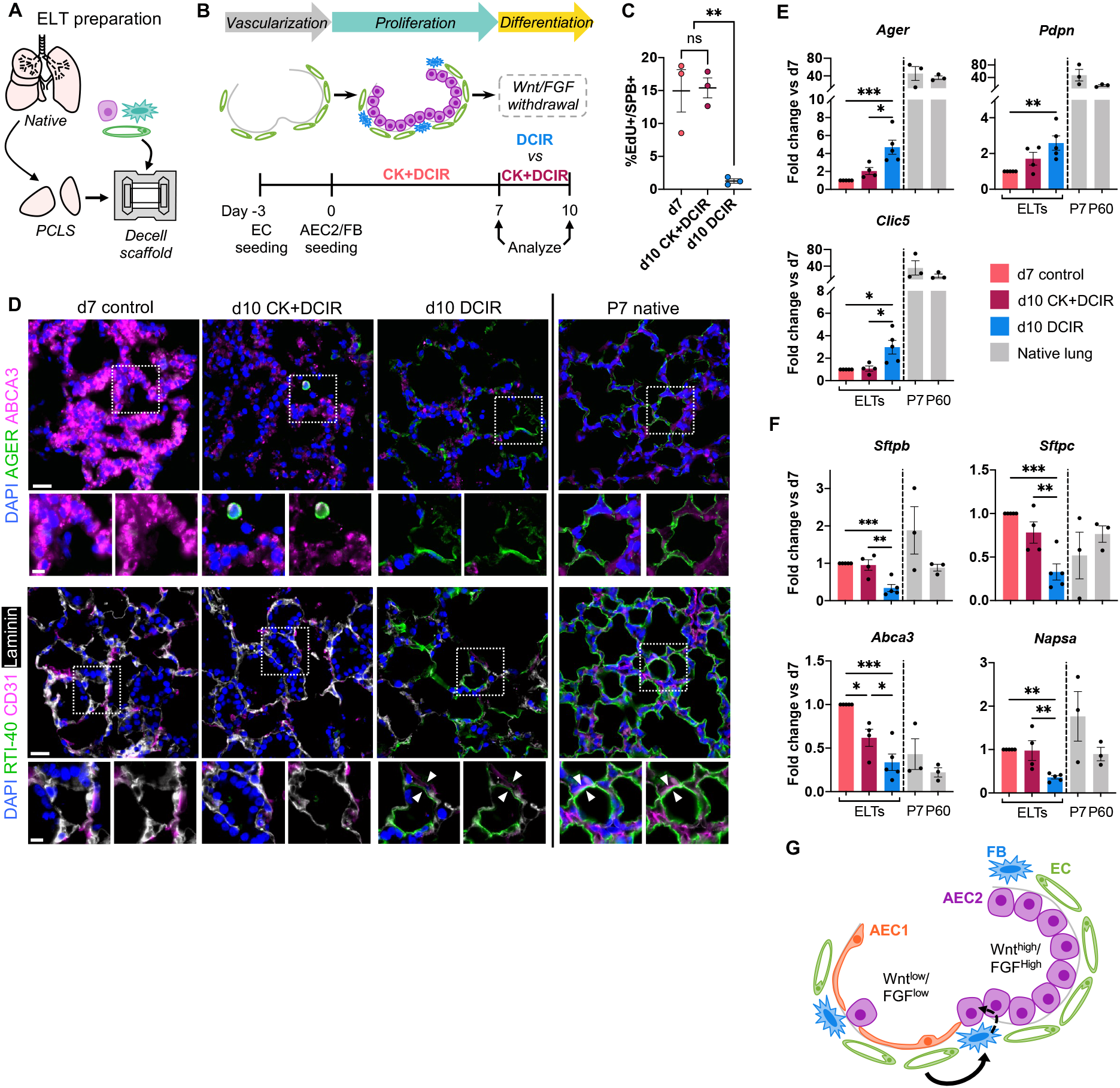
Wnt/FGF withdrawal is a modest cue for AEC2 differentiation. (**A**) Schematic of tri-culture engineered lung tissue (ELT) preparation. (**B**) Timeline of experiment examining effects of CHIR and KGF removal on AEC2 differentiation in tri-culture ELTs. (**C**) Quantification of EdU^+^ AEC2s in ELTs via immunostaining (*n* = 3). (**D**) Immunostaining for AEC1 proteins AGER and RTI-40 in ELTs, with P7 native rat lung shown for comparison. Arrowheads indicate close apposition of an RTI-40^+^ AEC1 with a CD31^+^ EC. Scale bars: main image, 25 μm; magnified region, 10 μm. (**E** and **F**) qRT-PCR of AEC1 (E) and AEC2 (F) gene expression in ELTs. Native gene expression is normalized to the average expression in d7 tissues and shown for approximate comparison only. d10 CK+DCIR: *n* = 4; d7, d10 DCIR: *n* = 5; native: *n* = 3. (**G**) Proposed role for Wnt and FGF in regulating epithelial fate within the alveolus. (C, E, and F): Error bars: mean ± SEM. One-way ANOVA with Holm-Sidak. ns, not significant, \**P* < 0.05, \*\**P* < 0.01, \*\*\**P* < 0.001.

Engineered lung tissues cultured with CK+DCIR expressed AEC2 genes at near-native levels by day 7 of culture (Extended Data Fig. 4E). By immunostaining, tri-culture AEC2s expressed punctate pro-surfactant protein (SP)C and colocalized SPB/ABCA3, consistent with protein localization to lamellar bodies (Fig. 1E). Transmission electron microscopy (TEM) confirmed the presence of well-formed lamellar bodies in tri-culture AEC2s, as well as characteristic apical microvilli and tight junctions (Fig. 1F). There was also the presence of luminal tubular myelin, a structure characteristic of secreted surfactant (Fig. 1F). Airway fluid collected via bronchoalveolar lavage (BAL) of engineered lungs contained SPB at greater than 60% of native levels, as determined by enzyme-linked immunosorbent assay (Fig. 1G). This quantity of SPB is sufficient for normal physiologic lung function^28^.

To assess whole lung mechanical compliance, we generated pressure-volume curves. While decellularized scaffolds had a compliance that was significantly less than that of native lung, the compliance of engineered lungs upon initial expansion was not significantly different from that of native (Fig. 1H,I). This observation is consistent with the presence of functional surfactant in the engineered lungs, with a concomitant reduction in alveolar surface tension. Thus, engineered tri-culture lungs support the proliferation of abundant AEC2s within organized alveolar-like units (Fig. 1J). These AEC2s expressed native-like molecular and morphometric features^29^ and showed evidence of mechanically relevant surfactant secretion.

### Endothelial cells and fibroblasts are insufficient as independent niche cells

We next isolated the role of FBs within the engineered niche, by seeding lungs with AEC2s and ECs only, and omitting FBs from the culture system (Fig. 2A). Despite initial AEC2 engraftment not substantially different from that in FB-containing cultures, AEC2/EC lungs did not support AEC2 growth by day 7 (Fig. 2B-D, Extended Data Fig. 4A,B,C,E). Rather, these lungs were characterized by numerous ECs and, surprisingly, a number of FB-like cells (Fig. 2C,D, Extended Data Fig. 6A,B), despite the initial omission of FBs in the scaffold. In a separate experiment, we determined that these FB-like cells were derived from the primary AEC2 isolate (Extended Data Fig. 6C,D), and we speculate that they are most likely a primary cellular contaminant (Extended Data Fig. 1B,C). The initial scarcity of this mesenchymal contaminant (Extended Data Fig. 6B), which would have limited opportunities for epithelial cell contact, means these cells were likely unable to efficiently support AEC2 growth.

**Fig. 6.**
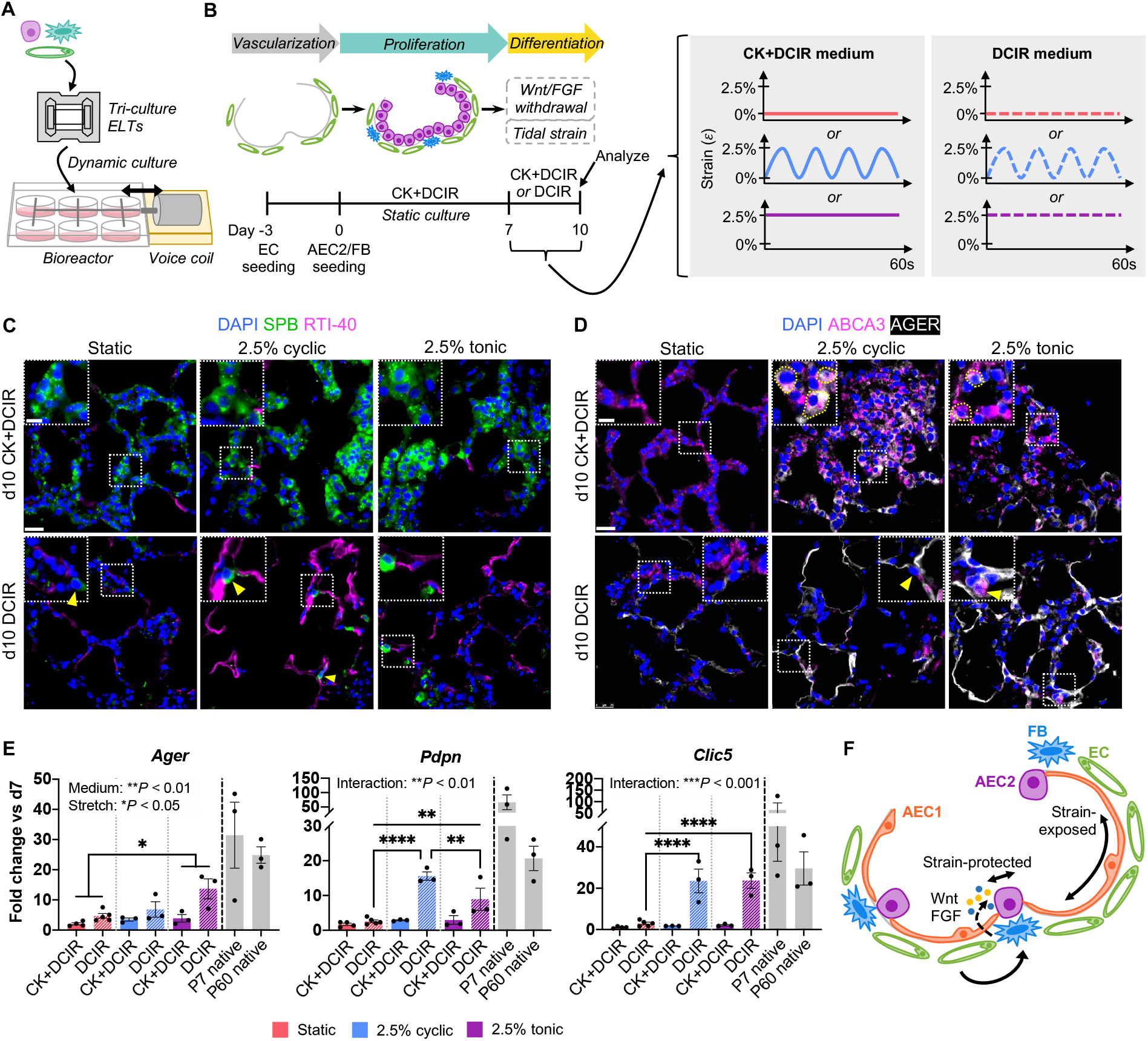
Tidal strain synergizes with biochemical cues to promote the AEC1 fate. (**A**) Schematic of bioreactor for engineered lung tissue (ELT) dynamic culture. (**B**) Timeline of experiment examining effects of CHIR and KGF withdrawal and tidal strain on AEC2 differentiation in tri-culture ELTs. (**C** and **D**) Immunostaining for AEC2 (SPB, ABCA3) and AEC1 (RTI-40, AGER) markers in tri-culture ELTs at d10. Yellow arrowheads, AEC2s localized to alveolar corners. Yellow dotted outlines, AGER^+^/ABCA3^+^ cells. Scale bars: main image, 25 μm; magnified region, 10 μm. (**E**) qRT-PCR of AEC1 gene expression in ELTs, normalized to their respective d7 control tissues. Native gene expression is normalized to the average expression in d7 tissues and shown for approximate comparison only. Static/CK+DCIR: *n* = 4; static/DCIR: *n* = 5; all other conditions: *n* = 3. Error bars: mean ± SEM. Text indicates significance by two-way ANOVA for medium main effect (“medium”), stretch main effect (“stretch”), and interaction between medium/stretch (“interaction”). *P* values for annotated pairwise comparisons were calculated via Holm-Sidak. \**P* < 0.05, \*\**P* < 0.01, \*\*\*\**P* < 0.0001. (**F**) Proposed role for Wnt/FGF and tidal strain in regulating epithelial fate within the alveolus.

Intriguingly, FBs in the absence of ECs were also insufficient as an independent niche cell. At day 7 of culture, AEC2/FB lungs contained disorganized epithelial clusters within a chaotic interstitium of FBs and collagenous matrix (Fig. 2E-G, Extended Data Fig. 6E,F): a tissue structure that is incompatible with gas exchange. Despite total and proliferating AEC2 numbers that were not significantly different from those in tri-culture lungs (Extended Data Fig. 4A-D), as well as similar whole-tissue AEC2 gene expression (Extended Data Fig. 4E), close inspection revealed only weak surfactant protein expression, without consistent punctate localization, in AEC2/FB co-cultured AEC2s (Fig. 2H). TEM demonstrated aberrant epithelial cell ultrastructure, including enlarged nuclei and irregular apical projections (Fig. 2I). These features suggest a loss of the appropriate AEC2 cellular program in the setting of FB-only co-culture in lung scaffolds. Indeed, we observed abnormal protein expression of alpha smooth muscle actin (αSMA) and procollagen Iα1 in the AEC2/FB epithelium (Extended Data Fig. 6G). AEC2/FB lungs also had epithelial and mesenchymal expression of the pro-fibrotic cytokine transforming growth factor beta 1 (TGFβ1) and its key transcriptional regulator phospho-SMAD3 (pSMAD3) (Fig. 2J), suggesting that AEC2/FB lungs display a pro-fibrotic milieu, with associated loss of AEC2 phenotype and disruption of cultured alveolar structure (Fig. 2K). Together, these data underscore the importance of an initial seeded FB population to support AEC2 growth in engineered lung, but suggest that the endothelium is critical for maintenance of AEC2 phenotype and proper alveolar organization in the presence of FBs (Fig. 1J).

### Endothelial cells modulate fibroblast niche support

We used single-cell RNA sequencing (scRNAseq) of day 7 AEC2/FB (“co-culture”) and tri-culture (AEC2/FB/EC) engineered lungs to further investigate the changes introduced to the alveolar niche by the presence of endothelium (Extended Data Fig. 7A-C, Supplementary Table 1). (Importantly, the endothelium in tri-culture lungs most resembles native lung capillary endothelium^30^; Extended Data Fig. 7D,E). We mapped a known database of ligand-receptor (LR) pairs to the scRNAseq data in order to characterize putative signaling networks within each engineered lung^31^ (Fig. 3A, Supplementary Table 2). This analysis suggested that ECs are involved in numerous paracrine signaling exchanges with both the epithelium and the mesenchyme in the tri-culture setting. We found that FBs, in both conditions, secrete known AEC2-supportive niche cues, such as *Wnt5a* and *Il6*^11, 18, 32^ (Fig. 3A,B). However, there was also expression of a number of unique signaling molecules that distinguished the culture conditions with and without endothelium, including prominent pro-fibrotic cytokine expression (*Ctgf*, *Tgfb1*) by co-culture FBs (Fig. 3B). Examination of key signaling pathways revealed that the endothelium introduced new signals into the engineered system (*e.g.* Notch signaling), but also fundamentally altered the epithelial-mesenchymal communication patterns, including abrogating (BMP/TGFβ) or shifting (PDGF) vectors of signaling (Fig. 3C). These data suggest that the role of the endothelium within the engineered tri-culture niche may include indirect effects on AEC2s, which are mediated via the FBs.

Thus, we utilized scRNAseq to further investigate FB phenotypic alterations in the presence of endothelium (Fig. 4A, Supplementary Table 3). FBs were cultured on polystyrene for 72 hours prior to lung seeding, and hence exhibited an overall shift toward a pro-migratory and contractile phenotype before culture in engineered lung scaffolds^33^ (Extended Data Fig. 8A). Comparison of differentially expressed genes among the cultured FB isolate, engineered lung FBs, and native lung FBs revealed that the polystyrene-expanded FB isolate was most different from native. Epithelial co-culture in lung scaffolds, and more so epithelial-endothelial tri-culture, caused shifts of previously polystyrene-cultured FBs back toward native FB phenotypes, despite a loss of population heterogeneity (Fig. 4A,B, Extended Data Fig. 8B).

Under conditions of AEC2/FB co-culture, but not tri-culture, we found that FBs retained elevated expression of *Acta2*, *Myl9*, and other contractile markers, as seen in the cultured FB isolate (Fig. 4C, Extended Data Fig. 8C,D). Co-culture FBs also uniquely upregulated pathways (mTORC1 signaling, reactive oxygen species [ROS] metabolism) and ECM-related genes (*Tgfb1*, *Col5a1*, *Col12a1*) that characterize FBs in pulmonary fibrosis^34, 35^ (Fig. 4C,D). Co-culture FB activation, as quantified by an unbiased activation score^35^, was modestly correlated (*r* = 0.64) with TGFβ/Smad3 activation (Fig. 4E, Extended Data Fig. 8E).

In contrast, ECM genes that were upregulated in tri-culture FBs included proteoglycans that are critical for proper collagen fibril assembly (*Dcn*, *Lum*) as well as a basement membrane collagen (*Col6a2*) that is critical for normal alveolar formation and AEC2 growth during alveologenesis^36^ (Fig. 4D). FBs in the presence of ECs in tri-culture lung were also enriched for genes associated with angiogenesis and lipid handling (Fig. 4D), and they upregulated signatures of mesenchymal subsets that have been proposed as specific AEC2 niche cells: *i.e. Axin2*^+^/*Pdgfra*^+^ mesenchymal alveolar niche cells (MANCs)^18^ and lipofibroblasts^37^ (Fig. 4E, Extended Data Fig. 8F,G). Together, these data imply that ECs modulate FB phenotype to maintain FB quiescence and to promote mesenchymal phenotypes that support the AEC2 niche (Fig. 4F).

### Wnt and FGF withdrawal is a modest cue for AEC1 differentiation

A mature alveolus requires squamous AEC1s for efficient gas exchange. However, in the studies described above, both co- and tri-culture engineered lungs demonstrated only minimal evidence of AEC2 differentiation to AEC1s by day 7, as assessed by immunostaining and qRT-PCR of engineered lung tissue (Extended Data Fig. 9). This is consistent with prior evidence suggesting that CK+DCI supplementation supports long-term maintenance of human pluripotent stem cell (hPSC)-derived AEC2s but does not support their further differentiation into AEC1s^23^. We therefore hypothesized that removal of Wnt agonist CHIR and FGFR2 agonist KGF (“CK”) on day 7 of culture, following epithelial expansion, would remove key signals favoring AEC2 proliferation and fate maintenance, and might permit their differentiation to AEC1s^11, 23, 32, 38^ (Extended Data Fig. 3D-F).

To test various combinations of cues for AEC1 differentiation within cultured lung scaffolds, we generated small-scale engineered lung tissues (ELTs) by reseeding acellular precision-cut lung matrices^39^ (Fig. 5A). ELTs were seeded with ECs followed by FBs and AEC2s, and cultured for an initial 7-day proliferation phase. Thereafter, we removed CK (“DCIR” medium) and maintained culture for an ensuing 3 days (“d10 DCIR”; Fig. 5B). Quantification of 5-ethynyl-2′-deoxyuridine (EdU) incorporation confirmed a significant decrease in AEC2 proliferation after 3 days of CK withdrawal (Fig. 5C). This was accompanied by the striking appearance of many DCIR-cultured engineered alveoli lined with squamous-appearing, AGER^+^ or RTI-40 (podoplanin)^+^ AEC1-like cells at day 10, with a microscopic tissue appearance somewhat approaching that of native lung (Fig. 5D). Some clusters of AEC2s remained (Fig. 5D). We also observed closely apposed RTI-40^+^ AEC1 and CD31^+^ EC cellular processes in day 10 DCIR tissues, mimicking the apposition that is characteristic of blood-gas barrier formation^27^ (Fig. 5D). qRT-PCR revealed a modest but significant increase in AEC1 gene expression, with a concomitant decrease in AEC2 gene expression, after the withdrawal of CK, as compared to day 7 ELTs (Fig. 5E,F). In contrast, ELTs that were cultured from days 7 to 10 with CK+DCIR (“d10 CK+DCIR”) exhibited no change in AEC2 proliferation, and no significant evidence of differentiation to AEC1s (Fig. 5C-F). This implies that the partial AEC1 differentiation that was observed with CK withdrawal was not simply a consequence of increased time in culture. Together, these data suggest that Wnt and FGFR2 agonism, via the CK factors, promotes the maintenance of AEC2 fate, and that CK withdrawal can induce some partial AEC1 differentiation (Fig. 5G).

### Tidal strain synergizes with Wnt/FGF withdrawal to promote the AEC1 fate

Fetal breathing movements are essential for AEC1 differentiation during lung development^40^, and accumulating evidence points to activation of the mechanosensitive YAP/TAZ pathway as requisite for the AEC1 fate^41, 42^. These observations imply a role for native-like mechanical forces in regulating alveolar epithelial differentiation. Thus, we tested the application of tidal-level mechanical strain as an alternative cue for AEC2 differentiation to AEC1s in engineered alveoli. Tri-lineage ELTs were repopulated with AEC2s via static culture through day 7 in the presence of CK+DCIR medium. ELTs were then transferred to a bioreactor^43^ for 3 additional days under either 2.5% cyclic or tonic uniaxial strain, in the presence of either CK+DCIR or DCIR medium (Fig. 6A,B). ELTs that were previously cultured through day 10 under static conditions served as no-strain controls.

After 3 days of strain, ELTs cultured with CK+DCIR through day 10 contained epithelial clusters and alveolar-like structures that were lined with cuboidal AEC2 monolayers (Fig. 6C,D, Extended Data Fig. 10B,C). Only a few flattened cells were observed that expressed AEC1 markers RTI-40 or AGER (Fig. 6C,D), and AEC1 gene expression by whole-tissue qRT-PCR was very low (Fig. 6E). However, tissues cultured with CK+DCIR under strain contained multiple cuboidal AGER^+^/ABCA3^+^ cells (Fig. 6D), which may represent AEC2s in an intermediate state of differentiation.

In contrast, DCIR-treated, stretched ELTs comprised alveolar-like structures that were lined by squamous AEC1s (Fig. 6C,D). RTI-40 and AGER protein expression was noticeably brighter and more widespread in stretched tissues than in tissues that were cultured statically with DCIR (Fig. 6C,D). Although AEC2 gene expression was significantly reduced following the removal of CHIR and KGF for the final 3 days of culture (Extended Data Fig. 10D), scattered AEC2s remained that were often localized to the alveolar corners, which is their typical location in the native lung^44^ (Fig. 6C,D, Extended Data Fig. 10A-C). qRT-PCR demonstrated a significant synergy between stretch and Wnt/FGF withdrawal in promoting AEC1 differentiation, with AEC1 gene expression approaching native levels in DCIR-treated, stretched tissues (Fig. 6E). The effect of cyclic stretch versus tonic stretch appeared to vary by marker, which suggests that the stretch pattern may confer differences in cell function or molecular state that are not captured by the analysis performed here (Fig. 6C-E). Taken together, these findings suggest that a Wnt- and FGF-low environment, coupled with tidal-level strain, promotes AEC2 differentiation to AEC1s within the engineered alveolus (Fig. 6F).

## DISCUSSION

The alveolus comprises a complex niche of factors that act in concert to support distal lung stem cell function and, ultimately, stable gas exchange. Here, we utilized a tissue engineering strategy to temporally modulate selected cellular, biochemical, and mechanical niche cues to generate epithelial-mesenchymal-endothelial alveolar units. These studies take a pivotal step toward the formation of a population of functional alveoli that may one day be capable of gas exchange in engineered lung. Importantly, our biomimetic engineered systems also uniquely allow the parsing of impacts of diverse niche constituents that cannot be successfully teased apart in complex living systems. The lessons learned from these engineered lung tissues may pave the way for direct experimental confirmation of key alveolar cues *in vivo*, and add to our understanding of alveolar formation and maintenance.

The nascent engineered alveoli in this report comprise three distinct cell lineages and contain both AEC2s and AEC1s having native-like phenotype and organization relative to other cell types. These engineered alveoli have substantially more fidelity to native structures than previous reports of organoid cultures, or any other reported engineered lung systems. Although our studies used neonatal AEC2s, whose fate regulation may differ from that of adult AEC2s^38, 42^, this work provides proof of principal for a “directed regeneration” approach, whereby tissue repair by delivered cells might be deliberately induced via careful modulation of the tissue microenvironment. Our strategy also circumvents the challenges associated with isolating primary AEC1s or differentiating AEC1s at scale from hPSCs for tissue repopulation^23, 45^, by instead deriving AEC1s *in situ* from seeded AEC2s within a spatially complex organ scaffold.

This work highlights the diversity of cues that coordinate epithelial cell fate within the alveolus, and suggests caution in drawing conclusions about stimuli that are tested in isolation. Our proliferation experiments suggest a key role for ECs in maintaining FB quiescence and in promoting mesenchymal AEC2 niche support. These studies augment prior evidence that ECs can support epithelial repair^20, 46^ and patterning^47^ via crosstalk with the mesenchyme. Our finding that tidal-level strain synergizes with Wnt/FGF withdrawal to induce AEC1 differentiation sheds additional light on previous observations that neither Wnt inhibition^45^ nor genetic YAP activation in AEC2s (in effect, activating some of the downstream signaling induced by mechanical stretch, via molecular tools)^41, 42^ is sufficient to induce full differentiation to squamous AEC1s. It may be that local variation in strain levels within the alveolus^48^ acts in tandem with short-range FB-derived Wnts^11^ to determine epithelial differentiation state.

The current study, by focusing on recapitulating cellular-level epithelial phenotypes *in vitro*, addresses an important stumbling block in the path toward regenerating functional lung tissue. Additional functional assessments and evaluation of long-term tissue durability will be essential. Integrating the strategies presented here with previous advances in bioengineered lung microvascular regeneration^5, 6^, and incorporating yet additional niche complexity, including alveolar macrophages and an air-liquid interface, will likely be additional important steps toward improving engineered lung phenotype.

## METHODS

### Animals

All animal work was performed in accordance with AAALAC guidelines and was approved by the Yale Institutional Animal Care and Use Committee. Animal husbandry and veterinary care were provided by the Yale Animal Resources Center. Animals were group housed, provided clean bedding and *ad libitum* water and food, and housed under controlled temperature and humidity with a 12 hr light/dark cycle. Wild-type Sprague-Dawley rats were purchased from Charles River Laboratories and used for decellularized lung scaffold preparation and primary AEC2 and fibroblast isolations. GFP^+^ AEC2s were isolated from the lungs of SD-Tg(CAG-*EGFP*)4Osb (*EGFP*^+^) rats, in which *EGFP* is driven by the ubiquitous CAG promoter (originally from Dr. Masaru Okabe, Osaka University, Japan)^49^.

### Isolation of rat AEC2s

Rat alveolar epithelial type 2 cells (AEC2s) were isolated from 7-9-day-old wild-type or *EGFP*^+^ Sprague-Dawley rat pups as previously described^50^, with slight modifications. For each isolation, 5-10 pups were randomly selected from a single litter without bias for males or females. For a given experiment, each biological replicate used an independent AEC2 isolate. Briefly, pups were euthanized via intraperitoneal (IP) injection of sodium pentobarbital (150 mg/kg). The thorax was entered and the lungs were flushed of blood by intracardiac injection of 100 U/mL heparin (Sigma) and 0.01 mg/mL sodium nitroprusside (Sigma) in PBS. The trachea was cannulated with a 24 G, 0.75-inch intravenous catheter (BD). The lungs were inflated with dispase (2 U/mL; Roche) in Dulbecco’s modified Eagle’s medium (DMEM; Gibco) with 20% HEPES-buffered saline (50 mM HEPES [Corning] and 150 mM NaCl, pH 7.4 in dH_2_O), followed by 1% low-melting-point (LMP) agarose (Invitrogen). The agarose was gelled under ice, and then the lungs were extracted and incubated in additional dispase solution for 45 min at room temperature. The distal lung tissue was separated from the mainstem bronchi and transferred to a Petri dish containing DMEM with 1% HEPES and 100 U/mL DNase I (Worthington). The tissue was then dissociated with a blunt bone marrow aspiration needle, 1 mM EDTA (AmericanBio) was added, and the cell suspension was sequentially filtered through 100-(Fisherbrand) and 40-μm (Falcon) strainers, and 20-μm nylon mesh (Millipore). The cells were labeled with RTII-70 antibody (Terrace Biotech)^51^ in buffer (Hank’s balanced salt solution [HBSS; Gibco] with 2mM EDTA and 0.5% BSA [Gemini Bio]), tagged with anti-mouse IgG microbeads (Miltenyi), and magnetically sorted via MS columns (Miltenyi). Freshly isolated cells were used immediately in all experiments, without intervening culture.

### Isolation and culture of rat lung fibroblasts

Neonatal rat lung fibroblasts (FBs) were isolated from 7-9-day-old Sprague-Dawley rat pups using a method similar to that described by Bruce and Honaker^52^. For each isolation, 5-10 pups were randomly selected from a single litter without bias for males or females. For whole lung experiments, each replicate used cells from an independent FB isolation. Pups were euthanized, the chest was entered, and lungs were flushed of blood as described for the isolation of AEC2s and excised *en bloc*. Following removal of the heart and hilar structures, the lungs were minced and digested in 0.5 mg/mL trypsin (Gibco) with 0.3 mg/mL collagenase type I (Worthington) in HBSS for 2 hours in a shaking water bath at 37°C, with trituration every 15 min. One hour into the digestion, the cells in suspension were removed and mixed 1:1 with DMEM containing 10% FBS (Hyclone). Fresh enzyme solution with 100 U/mL DNase I was added to the remaining tissue pieces, and incubation continued for 1 hr. At the end of the incubation, any remaining tissue pieces were dissociated with a blunt bone marrow aspiration needle and mixed 1:1 with DMEM with serum. The combined cell suspensions were filtered through 100- and 70-μm (Falcon) strainers, centrifuged at 400 g for 10 min at 4°C, and resuspended in HBSS. The cells were plated at 30-40 million cells per T75 cell culture flask with the balance in volume to 15 mL per flask made up by DMEM with 10% FBS and 1% penicillin/streptomycin (P/S [Gibco], final serum concentration 8%). The flasks were incubated at 37°C for 60 minutes, then aspirated of non-adherent cells, rinsed with PBS, and replenished with fresh medium (DMEM + 10% FBS + 1% P/S). On day 1 after plating, the flasks were rinsed once more with PBS and replenished with fresh culture medium. For whole lung cultures, freshly isolated FBs were cultured approximately 72 hours in a humidified incubator at 37°C and 5% CO2, with a medium change after 24 hrs, and used at passage 1. For alveolosphere and ELT (slice) cultures, FBs were used at passage 1-2.

### Endothelial cell culture

Primary rat lung microvascular endothelial cells (RLMVEC, VEC Technologies) were cultured on fibronectin (1 μg/cm^2^; Sigma)-coated flasks in MCDB-131 Complete medium with serum (VEC Technologies) and used at passage 5-6. Cells were cultured in a humidified incubator at 37°C and 5% CO2, with medium changes every other day.

### Whole lung decellularization

Decellularized whole lung scaffolds were prepared from the lungs of 300-350 g male Sprague-Dawley rats (approximately 8-12 weeks of age). Animals were anesthetized via IP injection of ketamine (75 mg/kg) and xylazine (5mg/kg), followed by IP injection of heparin (400 U/kg). The thorax was entered and lungs were cannulated *in situ* via the trachea and pulmonary artery (PA) with 1/16-inch barbed fittings (Cole Parmer), then cleared of blood via PA perfusion with 100 U/mL heparin and 0.01 mg/mL sodium nitroprusside in PBS with simultaneous manual ventilation of the airways via 10 mL syringe. Lungs and heart were extracted *en bloc* and decellularized with a Triton X-100/sodium deoxycholate (SDC)-based protocol as previously described^24^. Briefly, lungs were rinsed with antibiotics/antimycotics (10% P/S, 4% amphotericin-B [Sigma], 0.4% gentamicin [Gemini Bio] in PBS with Ca^2+^ and Mg^2+^ [Gibco]) followed by perfusion with 0.0035% Triton X-100 (American Bioanalytical). Lungs were inflated via the trachea with benzonase endonuclease (20 U/mL; Sigma) and incubated for 30 min, then rinsed via the PA with 1M NaCl and a gradient of sodium deoxycholate solutions (0.01%, 0.05%, 0.1%; Sigma). Next, lungs were perfused with 20 U/mL benzonase and incubated for 60 min. Finally, lungs were perfused with 0.5% Triton X-100 and rinsed extensively with PBS. Decellularized scaffolds were stored in antibiotics/antimycotics solution for up to 30 days at 4°C before use.

### Engineered whole lung culture and harvest

For engineered whole lung cultures, lung scaffolds were mounted in our previously described bioreactor system with oxygenation via hollow-fiber cartridge^26^. For endothelial cell-seeded lungs, scaffolds were pre-seeded on day -3 of culture with 38.0 ± 1.4 M RLMVEC via the pulmonary veins (PV) and 38.8 ± 1.4 M RLMVEC via the PA as previously described^25^, and cultured for 3 days in MCDB-131 complete medium with 2% FBS, 1% P/S, and 0.1% gentamicin with 20 mL/min perfusion via the PA, in an incubator at 37°C with 5% CO2. On day 0 of culture, the bioreactor medium was replaced with CK+DCIR medium and scaffolds were seeded with 40 M AEC2s with or without 36.4 ± 1.8 M FBs via injection into the trachea. Following both endothelial and epithelial cell seedings, the lung was allowed to sit statically for 90 min at 37°C. PA perfusion was then resumed at 2 mL/min, and ramped up to 20 mL/min over the course of 3 hours. CK+DCIR medium consists of epithelial base medium (50% DMEM [Gibco], 50% F-12 Nutrient Mixture [Gibco], 15 mM HEPES [Corning], 4 mM L-glutamine [Gibco)] 1% P/S [Gibco], 0.1% gentamicin [Gemini Bio], 10 μg/mL insulin [Sigma], 5 μg/mL transferrin [Sigma], 0.1% BSA Fraction V [Gemini Bio]) supplemented with CK+DCIR additives (3 μM CHIR99021 [CHIR; PeproTech], 10 ng/mL keratinocyte growth factor [KGF; PeproTech], 50 nM dexamethasone [Sigma], 0.1 mM 8-Bromo cAMP [Sigma], 0.1mM 3-Isobutyl-1-methylxanthine [IBMX; Sigma], 0.01 μM retinoic acid [Sigma]) (adapted from refs.^23, 53^). On day 3 of culture following epithelial seeding, the right lower lobe was tied off and removed. The lobe was divided for qRT-PCR analysis (snap frozen) and histological analysis (injected to inflate with 10% neutral-buffered formalin [NBF] and fixed for 4 hours). Lung PA perfusion was decreased to 14 mL/min and culture continued for an additional 4 days. Throughout culture, partial media changes were performed daily, titrating to maintain medium glucose > 70 mg/dL and lactate < 10 mM (monitored using glucose test strips [GlucCell] and lactate test strips [Nova Biomedical]). The remaining engineered lung lobes were harvested on day 7 of culture following epithelial seeding. Lobes were either snap frozen; inflated with 10% NBF at 10-15 cm cmH_2_O via the trachea, then allowed to fix for 4 hours prior histological processing; or dissociated for scRNAseq (see details below).

### Whole lung assays

Native lungs were extracted from 300-400 g male rats as described for decellularized lung scaffold preparation. Decellularized lungs were generated as described above. The right lower lobe of native and decellularized lungs was tied off in order to match day 7 engineered lungs.

#### Pressure-volume curves

For engineered lungs, quasi-static pressure-volume (PV) curves were generated on day 7, prior to lobe freezing or fixation. Lungs were connected via trachea cannula to a syringe fitted with a pressure transducer (Edwards), with real-time monitoring in LabChart 8 (ADInstruments), then inflated three times with air to total lung capacity. To generate PV curves, lungs were then inflated stepwise to 1 mL total volume, followed by stepwise deflation to resting volume, with a 15 sec pause after each 0.1 mL volume change. Pressure measurements were captured from the end of each 15 sec interval, and plotted against lung volume. Three PV curves were generated for each lung. Quasi-static compliance was calculated from the slope of the opening limb of each PV curve, and is reported as the average of three technical replicates per lung.

#### Bronchoalveolar lavage

For engineered lungs, bronchoalveolar lavage (BAL) was performed on day 7 after PV curve testing. One volume of PBS (equal to 0.7 x 30 mL/kg rat weight) was gently inflated and suctioned via the trachea, collected on ice, and spun down to remove cells. BAL was analyzed by ELISA for SPB protein (LSBio) according to the manufacturer’s instructions.

### 3D lung alveolosphere assays

We modified the previously described alveolosphere assay^54^ to take advantage of the tri-culture, serum-free system described in this study. Co- and tri-culture alveolosphere cultures were prepared in 0.4 μm, 24-well format cell culture inserts (Falcon) with 1:1 growth-factor reduced Matrigel (Corning) to cell suspension in epithelium base medium (90 μL total volume, 2.25 x 10^4^ AEC2s, 2.25 x 10^4^ FBs, and/or 4.5 x 10^4^ RLMVECs per insert; the number of AEC2s was kept constant across all co- and tri-culture conditions). Matrigel was allowed to solidify at 37°C for 30 min, and then 0.5 mL medium was added to the outside of each insert. Culture media were as follows: Extended Data Fig. 3A-C: CK+DCIR or conditioned medium (CM, see “Preparation of conditioned medium” below), Extended Data Fig. 3D-F: CK+DCIR or CK+DCIR with the specified medium component(s) removed (*e.g.* CK+DCIR without CHIR = “-CHIR”). Culture medium was changed on day 1 and every 2 days thereafter, and alveolospheres were analyzed after 7 days. 2-3 wells per condition were analyzed for each biological replicate, with reported quantification values (see “Image quantification” below) representing the average of technical replicate wells. Alveolospheres were imaged with an Axio Vert.A1 inverted microscope (Zeiss) and a Lumenera camera.

### Preparation of conditioned medium

To prepare conditioned medium (CM) for the 3D alveolosphere assay, freshly isolated FBs or passage 4-5 RLMVECs were cultured to 70-80% confluence in growth medium, then rinsed twice with 50/50 DMEM/F12 before exchanging the medium for CK+DCIR. Cells were cultured for 24 hrs more, at which point the CM was collected, sterile filtered, and stored at -80°C until use. Endothelial CM was used undiluted. Fibroblast CM was used diluted 1:1 with fresh CK+DCIR medium, as preliminary experiments revealed significant apoptosis in the setting of undiluted FB-CM.

### Engineered lung (slice) tissue (ELT) preparation and culture

ELTs were prepared as previously described^39^ and briefly outlined below.

#### Preparation of precision-cut lung slices

Native lungs were extracted from 300-400 g male rats as described for decellularized whole lung scaffold preparation, then inflated via the trachea with 2% LMP agarose in phenol red-free HBSS. The trachea was capped and the lung chilled on ice to allow the agarose to gel. The lobes were cut using a vibratome (Compresstome VF-300-0Z, Precisionary Instruments LLC) to 450 μm thick, and slices were snap frozen and stored at -80°C until use (not more than 3 months).

#### Slice decellularization

Lung slices were thawed under PBS, then cut into 3 mm-wide strips and clipped into polytetrafluoroethylene (PTFE) tissue-culture cassettes^55^. Slices were decellularized in 6-well plates using a Triton X-100/SDC-based protocol adapted from our previously described whole lung decellularization protocol (see above and ref.^24^). Decellularized slices were incubated with antibiotic/antimycotics for 48 hrs at 37°C, then stored at 4°C until seeding (not more than 30 days).

#### Slice seeding and culture

Tri-culture engineered lung (slice) tissues (ELTs) were prepared according to the same culture timeline and cell ratios used for tri-culture engineered whole lungs. Decellularized lung slices were rinsed with PBS, then transferred upside down to customized polydimethylsiloxane (PDMS) seeding baths^55^. Scaffolds were seeded on day -3 with ECs by pipetting a 100 μL cell suspension containing 5 M cells/mL of RLMVECs in endothelial medium (MCDB with 2% FBS, 1% P/S, and 0.1% gentamicin, 5 x 10^5^ cells per slice) directly onto the scaffold. After 2 hrs incubation at 37°C, 900 μL fresh medium was added to each well.

Endothelial medium was changed on day -2. On day 0, endothelial medium was removed, and the slices were seeded with a 100 μL cell suspension containing 2.5 M cells/mL AEC2s and 2.5 M cells/mL FBs in CK+DCIR medium (2.5 x 10^5^ AEC2s and 2.5 x 10^5^ FBs per slice). The slices were incubated at 37°C with 5% CO2 for 2 hrs, then 900 μL fresh CK+DCIR was added to each well. 24 hrs after seeding, the slices were removed from seeding baths, flipped, and transferred to well plates. The tissues were cultured in CK+DCIR medium until day 7, with medium changes every other day. On day 7, tissues were either saved for analysis or cultured for an additional 3 days (to day 10), either statically or under strain (see “Bioreactor Culture” below), and in either CK+DCIR or DCIR medium (*i.e.* with CHIR and KGF removed).

#### Bioreactor culture

ELTs were cultured under strain using an adaptation of our previously described bioreactor^43^. The bioreactor comprises a custom-made 3D-printed adapter (Formlabs Dental SG resin) which fits within a 6-well plate, allowing 6 ELTs to be stretched uniaxially in parallel. Each ELT is held by one fixed arm and one stainless-steel spring arm. The latter arm is driven by a voice coil linear actuator (BEI Kimco) controlled by a custom-programmed Pluto servo drive (Ingenia). Tissues were cultured statically in CK+DCIR medium through day 7, then removed from their PTFE frames and loaded into the bioreactor for 3 additional days of culture under either 2.5% cyclic (4 cycles/min) or 2.5% tonic uniaxial strain.

#### Takedown/analysis

For histological analysis and 5-ethynyl-2′-deoxyuridine (EdU) labeling, ELTs were incubated with 10 μM EdU for 2 hrs, then fixed with 10% NBF and paraffin embedded. For PCR analysis, RNA was extracted from fresh tissues and analyzed as described below; 2 ELTs were pooled per condition per experiment.

### Histology and immunofluorescent staining

Formalin-fixed tissues or Matrigel-embedded alveolospheres were routinely processed for paraffin embedding. Tissue sections were stained by standard methods for histological stains, or prepared for immunofluorescent (IF) staining. For IF, sections were baked for 45 min at 65°C, deparaffinized and rehydrated, and subjected to heat-mediated antigen retrieval with citric acid. Sections were permeabilized with 0.2% Triton X-100 in PBS for 10-15 min, blocked with blocking buffer (5% BSA and 0.75% glycine in PBS) for 1 hr at room temperature, and incubated with primary antibodies in blocking buffer overnight at 4°C. Sections were rinsed and secondary antibodies applied for 1 hr at room temperature, then stained with 4’,6-amidino-2-phenylindole (DAPI) and mounted with polyvinyl alcohol with DABCO (PVA-DABCO; Sigma). A list of antibodies and concentrations used for IF can be found in Supplementary Table 4. For EdU staining, sections were processed according to the manufacturer’s instructions (Click-iT™ EdU Cell Proliferation Kit for Imaging, Alexa Fluor™ 647, Invitrogen). Histological images were taken with an Axioskop 2 Plus upright microscope (Zeiss) and an AxioCam MRc camera (Zeiss). Fluorescent images were taken with a Leica DMI6000 B inverted fluorescent microscope and an Andor camera. Images were processed using ImageJ/Fiji. Any adjustments to brightness or contrast were applied equally across all experimental conditions in a given panel. In select instances, the contrast of individual color channels in a merged image was adjusted; this was performed identically across all experimental conditions.

### Oil Red O staining

Freshly isolated FBs were plated on a chamber slide (Lab-Tek II CC2, Nunc) and cultured for 72 hours, then fixed in 4% paraformaldehyde for 20 min at room temperature. Cells were stained with 0.18% weight/volume Oil Red O (Sigma) in 60% isopropanol for 30 min, then counterstained with hematoxylin for 1 min.

### Quantitative real-time PCR

Cell samples, whole engineered lung, ELTs, and native rat tissue samples were homogenized in lysis Buffer RLT (Qiagen). Total RNA was isolated using the RNeasy Mini Kit (Qiagen) following the manufacturer’s instructions. RNA was reverse transcribed using the iScript cDNA Synthesis Kit (Bio-Rad), according to the manufacturer’s protocol. All PCR reactions were run in triplicate using 1 μL of cDNA in a 25 μL final volume with iQ SYBR Green Supermix (Bio-Rad) and 200 nM each of forward and reverse primers (see Supplementary Table 5). qRT-PCR was performed on the CFX96 Real-Time System (Bio-Rad) for 40 cycles. Average threshold cycle values (Ct) from triplicate PCR reactions were normalized to *Actb* (β-actin) for whole engineered lungs, or to *B2m* for ELTs, and reported as fold change using the 2^-ΔΔCt^ method. Gene expression in engineered whole lung samples was normalized to the relevant starting cell population (*i.e.* AEC2 isolate for epithelial markers; FB isolate for mesenchymal markers); native P7 and P60 rat lung gene expression is normalized to that of the average starting cell population and is presented for approximate comparison only. Gene expression in ELTs was normalized to corresponding day 7 control tissues from the same experiment; native rat lung gene expression is normalized to that of the average day 7 control tissue and is presented for approximate comparison only.

### Transmission electron microscopy

Tissue was fixed for 30 min at room temperature followed by 90 min at 4°C in 2.5% glutaraldehyde and 2% paraformaldehyde in 0.1 M sodium cacodylate pH 7.4, and then rinsed 3 times in 0.1 M sodium cacodylate buffer. Processing and embedding of fixed tissue were performed by the Center for Cellular and Molecular Imaging, Electron Microscopy facility at Yale School of Medicine. Sections were imaged on a FEI Tecnai G2 Spirit BioTWIN Transmission Electron Microscope operated at 80 kV, using a Morada CCD camera (Olympus SIS).

### Sample preparation for scRNAseq

#### AEC2 and FB isolates

Freshly isolated AEC2s were resuspended in 0.01% BSA in PBS, filtered through a 40 μm cell strainer, and diluted to 1 x 10^6^ cells/mL for single-cell library preparation. Freshly isolated FBs were cultured for 72 hrs, trypsinized, resuspended in 0.01% BSA in PBS, filtered through a 40 μm cell strainer, and diluted to 1 x 10^6^ cells/mL for single-cell library preparation.

#### Native lung tissue

Six post-natal day (P)7 Sprague-Dawley rat pups (3 males and 3 females) were dissociated for scRNAseq as previously described, with modifications^56^. Pups were given a dose of 400 U/kg IP heparin and euthanized via IP injection of sodium pentobarbital (150 mg/kg). Lungs were cleared of blood as described for the isolation of AEC2s, cannulated with a 24 G catheter (BD), and then inflated via the trachea with 2-3 mL of pre-warmed (37°C) enzyme solution (1 mg/mL collagenase/dispase [Roche], 3 U/mL elastase [Worthington], 20 U/mL DNase [Worthington] in DMEM). The heart and large airways were dissected away and the remaining lung tissues and enzyme solution were collected and transferred to a shaking 37°C water bath at 75 rpm for 20 min. The tissue was passed through a wire mesh strainer and the strainer was rinsed with ice-cold DMEM + 10% FBS + 1% P/S. The tissue solution was centrifuged for 5 min at 300 g, resuspended in ACK lysing buffer (Lonza), and incubated for 90-120 sec to lyse red blood cells. The cells were diluted in 0.01% BSA in PBS, spun down, and passed through a 70 μm filter. The cells were spun again for 3 min at 300 g, resuspended in 0.01% BSA, then passed twice through 40 μm filters. The resulting single cell suspension was counted, assessed for viability, then serially diluted to ≤1 x 10^6^ cells/mL for single-cell library preparation.

#### Engineered whole lung

Approximately 1 cm^2^ of tissue from the right middle lobe of one AEC2/FB and one tri-culture engineered whole lung was dissociated for scRNAseq on day 7 of culture as described for P7 native rat lungs, with the following changes. Lung tissue was inflated with 5 mL of enzyme solution via repeated injection with a 25 G needle, then minced with fine scissors. The tissue and enzyme solution were collected, supplemented with an additional 5 mL of enzyme solution, and transferred to a shaking water bath as described above. No red blood cell lysis step was performed on engineered lung samples.

### Single-cell RNA sequencing and analysis

scRNAseq libraries were generated with the Chromium Single Cell 3’ v2 assay (for cell populations) or v3 assay (for native and engineered lung tissue), according to the manufacturer’s protocol (10x Genomics). Samples were diluted for an expected cell recovery of 5,000 cells (AEC2 and FB isolates) or 10,000 cells (all other samples). Libraries were sequenced on an Illumina HiSeq 4000 (for cell populations) or Illumina NovaSeq platform (for native and engineered lung tissue) with a targeted depth of 50,000 reads per cell. Alignment was performed using Cell Ranger 3.0.2 (10x Genomics), using Ensembl Rnor6.0 release 95.

#### Filtration, clustering, and embedding

Data was processed in R v3.6.1 using Seurat v3.1.1^57^. Cells were filtered to accept those having between 500-10,000 genes and between 500-100,000 UMI. Cells were additionally filtered based on the percentage of mitochondrial reads as follows: >1% and <7% for the AEC2 isolate; <5% for the FB isolate; <28% for P7 native rat lung; <25% for d7 AEC2/FB engineered lung; and >6% and <28% for d7 tri-culture engineered lung.

Expression matrices were normalized using the NormalizeData function and scaled with ScaleData, with regression on percentage mitochondrial reads, number of UMI counts, and cell cycle scoring. Each sample was clustered independently, and visualized using dimensionality reduction (UMAP, uniform manifold approximation and projection). As an additional filtration step, objects were subclustered by lineage (*Col3a1*^+^ mesenchymal cells [comprising fibroblasts, smooth muscle cells, pericytes, and mesothelium], *Epcam*^+^ epithelial cells, *Cdh5*^+^ endothelial cells, and *Ptprc*^+^ immune cells) to identify and remove suspected doublets or low-information cells. These filtered subsets were subsequently merged to create the full objects used for downstream analysis. Initial identities were designated by lineage assignment (see Supplementary Table 1 for cluster markers). Within the P7 native rat lung, these lineages were subsetted and further clustered to assign subpopulation identities based on known markers.

#### Differential gene expression analysis

Differential expression analysis was performed using FindAllMarkers in Seurat, using the default Wilcoxon Rank Sum test. Differentially expressed genes (DEGs) were defined as those genes expressed in a minimum of 10% of cells with fold change ≥1.5 and adjusted *P* < 0.05, unless otherwise specified. Gene ontological and pathway enrichment was performed using Enrichr^58^. Heatmaps were created using the package ComplexHeatmap^59^.

#### Cell scoring

Expression scoring for pre-defined gene sets was performed using the AddModuleScore function in Seurat (based on ref.^60^). Gene sets are listed in Supplementary Table 6 and were defined as follows: P7 endothelial subsets: top cluster markers among P7 endothelial populations; TGFβ1/Smad3 targets: ref.^61^; unbiased FB activation score: ref.^35^; P7 mesenchymal subsets: top cluster markers among P7 mesenchymal populations; MANC signature: ref.^18^. Lipofibroblast: curated from refs.^62, 63^.

#### Ligand-receptor analysis

The R package connectome v0.1.9^31^ was used to map gene expression against rat homologues of literature-supported and putative ligand-receptor (LR) pairs within the FANTOM5 database^64^. Preliminary filtering of LR pairs was based on expression in a minimum of 10% of cells and adjusted *P* < 0.05 for both ligand and receptor (see filtered connectomes in Supplementary Table 2). Edge weights, for visualization purposes, were calculated as the sum of the normalized expression values for ligand and receptor for a given ligand/source cell-target cell/receptor edge.

### Image quantification

Image quantification was not performed blinded, as the same investigator both performed the experiments and analyzed the data, and phenotypic differences were in many cases obvious between conditions.

#### Alveolospheres

For alveolosphere number and diameter quantification, two 5x brightfield images were taken from each of 2-3 wells per condition after 7 days of culture, and alveolospheres of at least 50 μm in diameter were manually counted and measured in Fiji^65^. Each replicate data point represents the average from 2-3 wells per condition.

#### Total and proliferating cell numbers

For quantification of total and proliferating AEC2s, or total FBs, in engineered whole lungs, 8 random 20x fields of view (FOV) were taken per sample (corresponding to 1,954-14,738 cells/sample), avoiding the large airways and vessels, and cells were counted in Fiji. For quantification of EdU^+^ AEC2s in immunostained ELTs, 7-8 random 40x FOV were taken per sample (corresponding to 429-1622 cells/sample), and cells were counted manually in Fiji.

### Statistical analysis

Statistical analysis was performed in Prism v9.0.0 (GraphPad). In all cases, data points represent biological replicates, and statistical tests were performed only on biological replicates (*n* ≥ 3). No sample-size calculations were performed. Other than preliminary filtering of single-cell RNAseq data, no data were excluded. In instances in which technical replicates were averaged to derive the presented data points, this information is specified in the corresponding methods section. The statistical tests used and exact value of *n* for each experiment are specified in the corresponding figure legend.

An unpaired two-tailed *t-*test was used for comparison of two groups. A one-way analysis of variance (ANOVA) followed by Holm-Sidak’s multiple comparisons test for planned pairwise comparisons was used for comparisons of more than two groups with one factor, with the following exception: alveolosphere assays were analyzed according to a randomized block experimental design (with each independent experiment as a block), with a repeated measures ANOVA for comparison of multiple groups, followed by Holm-Sidak’s multiple comparisons test. The conditioned medium alveolosphere assay had a missing value, as one replicate of the experiment did not include a FB-CM condition. Because a repeated measures ANOVA cannot handle missing values, we instead analyzed these data by fitting a mixed model as implemented in GraphPad Prism. This mixed model uses a compound symmetry covariance matrix, and is fit using Restricted Maximum Likelihood (REML). In the absence of missing values, this method gives the same *P* values and multiple comparisons tests as repeated measures ANOVA. In the presence of missing values (missing completely at random), the results can be interpreted like repeated measures ANOVA. Sphericity was assumed for randomized block experiments. For analysis of ELT experiments examining both stretch and medium factors, a two-way ANOVA with Holm-Sidak’s multiple comparisons test was used. For analysis of cell scoring using scRNAseq data, a Kruskal-Wallis test with Dunn’s post-tests was used to compare more than two groups.

For all tests, *P* < 0.05 was considered significant. For bar and scatter plots, center values represent the mean and error bars represent standard error of the mean (SEM). For boxplots, the center value represents the median; the limits of the box represent the 1^st^ and 3^rd^ quartile, respectively; and the whiskers extend to the minimum and maximum values. Outliers (defined as those data points greater than 1.5 times the interquartile range from the median) were removed from boxplots for visualization purposes only.

## DATA AVAILABILITY

The scRNAseq data generated in this study have been deposited in the Gene Expression Omnibus (GEO) database under accession code GSE178405. All data supporting the findings in this study are present within the article and associated supplementary information files.

Additional data related to this paper and code necessary to reproduce this work are available from the corresponding author upon reasonable request.

## Supporting information

Summary of supplemental tables, Supplementary Tables 4 and 5

## ACKNOWLEDGEMENTS

The authors thank the Niklason lab, E. Calle, and N. Chavkin for helpful discussions; and E. Calle and A. Cherskov for critical comments on the manuscript. We thank the staff of Yale Pathology Tissue Services, the Yale Center for Genome Analysis, and the Center for Cellular and Molecular Imaging, Electron Microscopy facility at Yale School of Medicine for assistance with the work presented here. This work was supported by NIH grants F30HL143880 (K.L.L.); the Medical Scientist Training Program Training Grant T32GM136651 (K.L.L. and M.S.B.R.); F30HL143906 (M.S.B.R.); HL146056 and EB017103 (K.K.H.); R01HL127349, R01HL141852, and UH2HL123886 (N.K.); U01HL145567 (N.K. and L.E.N.); R01HL152677 (E.L.H.); and R01HL138540 (L.E.N.). This work was also supported by AHA postdoctoral fellowship 20POST35210709 (Y.Y.), by NSF CAREER award 1653160 (S.G.C.), by a generous gift from Three Lakes Partners (N.K.), and by an unrestricted research gift from Humacyte Inc. (L.E.N.).

## AUTHOR CONTRIBUTIONS

Conceptualization: K.L.L., L.E.N. Data curation: K.L.L. Formal Analysis: K.L.L. Funding Acquisition: N.K., L.E.N. Investigation: K.L.L., Y.Y., T.S.A., P.B. Methodology: K.L.L., Y.Y., R.N., M.S.B.R., K.K.H., S.G.C., E.L.H., L.E.N. Resources: R.N., S.G.C., N.K. Software: K.L.L., M.S.B.R. Supervision: K.K.H., S.G.C., E.L.H., L.E.N. Visualization: K.L.L., L.E.N. Writing – original draft: K.L.L. Writing – review & editing: K.L.L., L.E.N.

---

**Extended Data Fig. 1.**
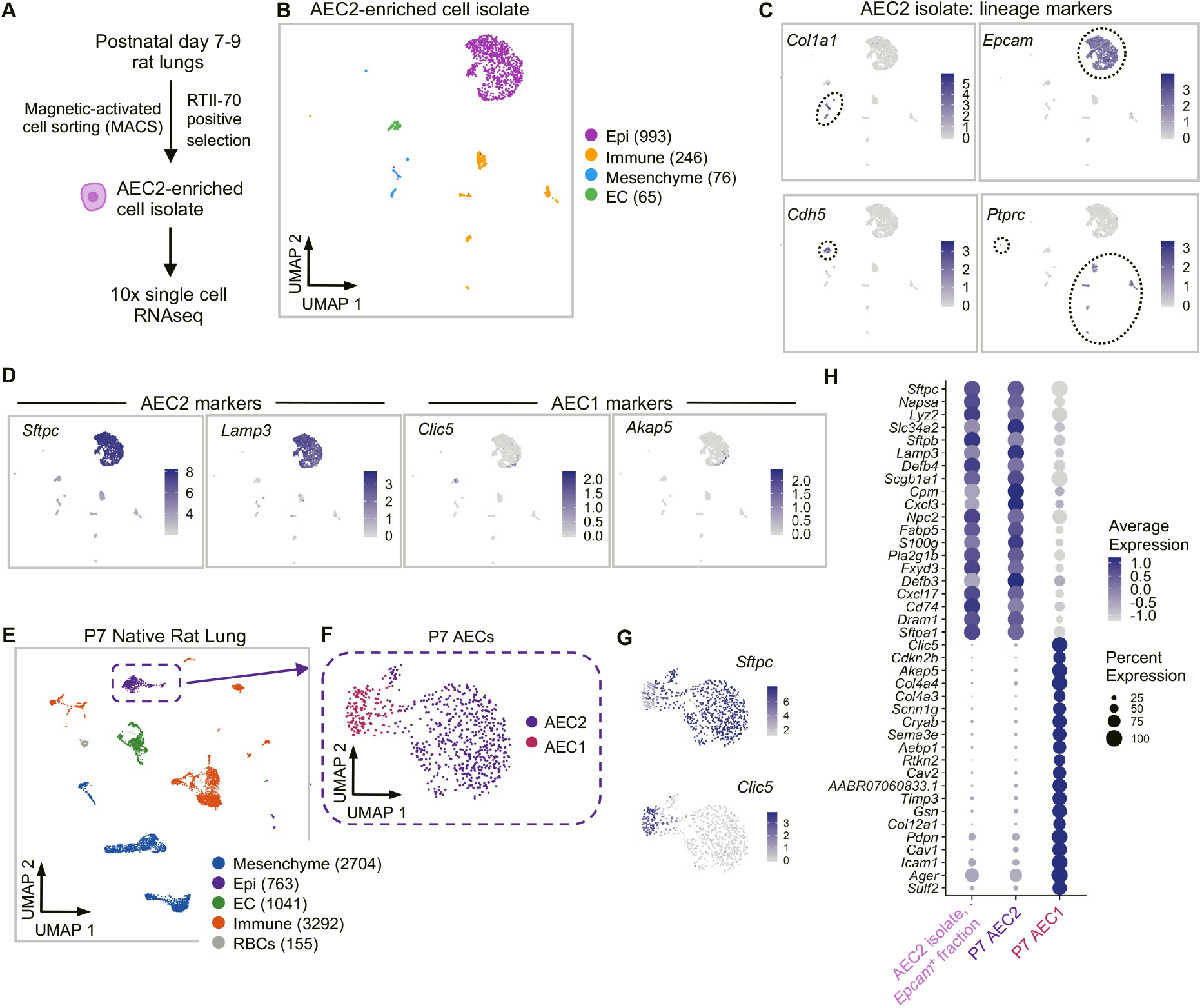
Characterization of the AEC2 isolate. (**A**) Strategy for isolation of rat AEC2s. (**B**) UMAP embedding of scRNAseq data for freshly isolated AEC2s. (**C** and **D**) Expression data for example markers used to assign cluster labels (C) and specific AEC2 and AEC1 markers (D), projected onto the scRNAseq data from (B). Dotted ovals encircle clusters positive for the indicated marker. (**E**) UMAP embedding of scRNAseq data for P7 whole rat lung dissociation. (**F**) UMAP embedding of native P7 AEC2s and AEC1s, extracted from the full P7 data in (E) and re-clustered. (G) Expression data for specific AEC2 (*Sftpc*) and AEC1 (*Clic5*) markers projected onto the scRNAseq data from (F). (**H**) Dot plot of gene expression for the top 20 P7 native AEC2 and AEC1 genes, demonstrating that the epithelial (*Epcam*^+^) fraction of the AEC2 isolate comprises AEC2s. (B and E): Clusters are colored by cell lineage. Numbers indicate cell numbers after filtering. Epi, epithelium. EC, endothelial cell. RBC, red blood cell.

**Extended Data Fig. 2.**
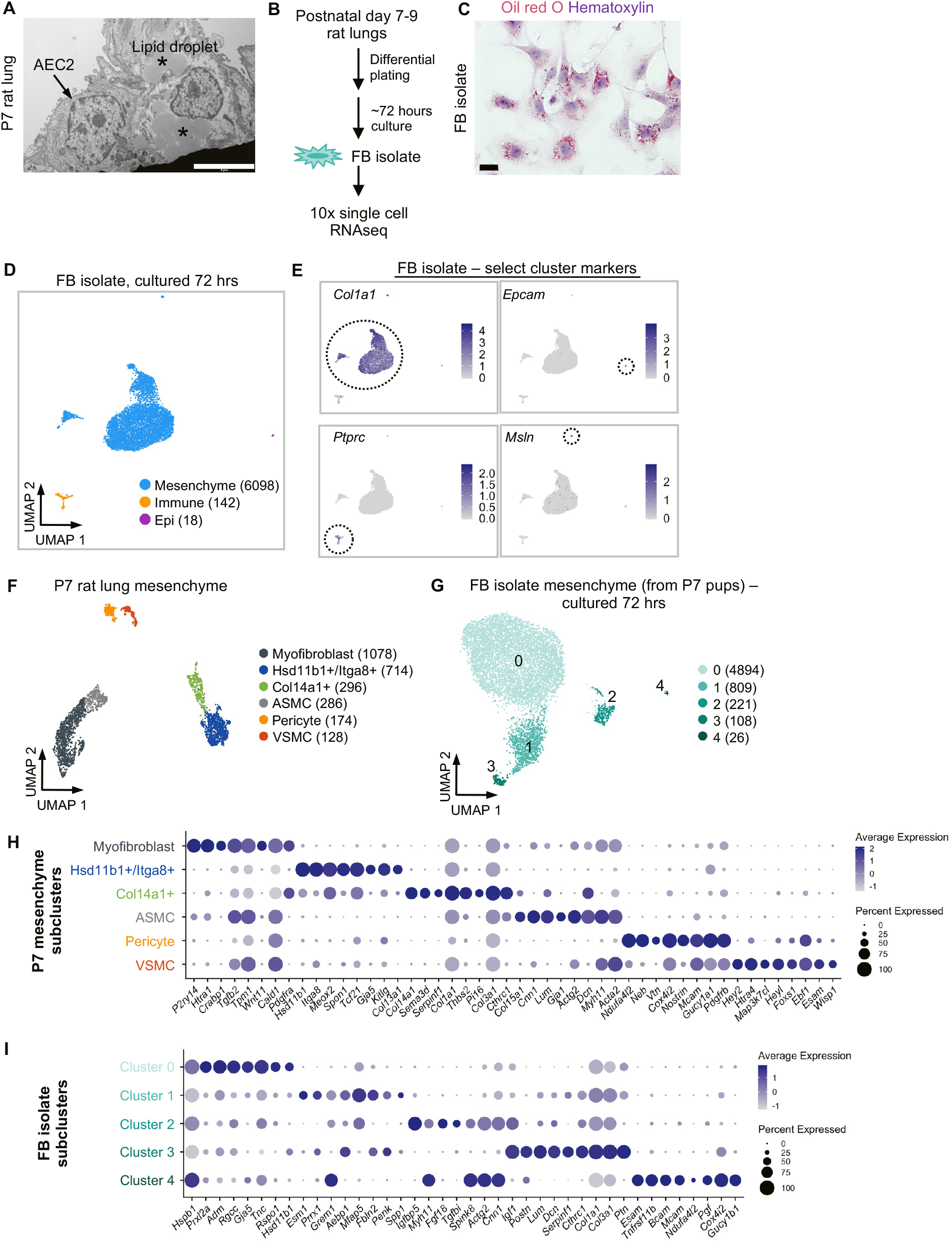
Preliminary characterization of the fibroblast isolate. (**A**) TEM of P7 rat lung demonstrating lipid-rich FBs neighboring an AEC2 at this timepoint. Asterisk, lipid droplet. Scale bar, 5 μm. (**B**) Strategy for isolating lipofibroblast-enriched neonatal rat lung fibroblasts (FBs). Cells were expanded on tissue-culture plastic for 72 hours following isolation in order to achieve sufficient numbers for whole lung scaffold reseeding. (**C**) Oil Red O staining of neutral lipid droplets in passage 0 FBs cultured for 72 hours. Scale bar, 20 μm. (**D**) UMAP embedding of scRNAseq data for isolated FBs cultured for 72 hours. Clusters are colored by cell lineage. Epi, epithelium. (**E**) Expression data for select lineage markers (*Col1a1*, *Epcam*, *Ptprc*) and mesothelial marker *Msln* projected onto the scRNAseq data from (D). *Msln*+ mesothelial cells were excluded from downstream analyses of FB isolate mesenchymal clusters. Dotted ovals encircle clusters positive for the indicated marker. (**F**) UMAP embedding of native P7 rat lung mesenchyme (excluding mesothelial cells), extracted from the full P7 data in Extended Data Fig. 1E and re-clustered. ASMC, airway smooth muscle cell. VSMC, vascular smooth muscle cell. (**G**) UMAP embedding of scRNAseq data for FB isolate mesenchymal cells (excluding mesothelial cells), extracted from the full FB isolate data in (D) and re-clustered. (**H**) Dot plot showing scaled expression of select top markers of P7 native lung mesenchyme clusters from (F). (**I**) Dot plot showing scaled expression of select top markers in FB isolate clusters from (G). Note that following 72 hours of culture on tissue culture plastic, the FB isolate preserved features of native heterogeneity, including subpopulations sharing characteristic markers of P7 native Hsd11b1^+^/Itga8^+^ FBs (Cluster 0), Col14a1^+^ FBs (Clusters 1 and 3), myofibroblasts and ASMCs (Cluster 2), and pericytes and VSMCs (Cluster 4). (D, F, and G): Numbers in parentheses indicate cell numbers per cluster after filtering.

**Extended Data Fig. 3.**
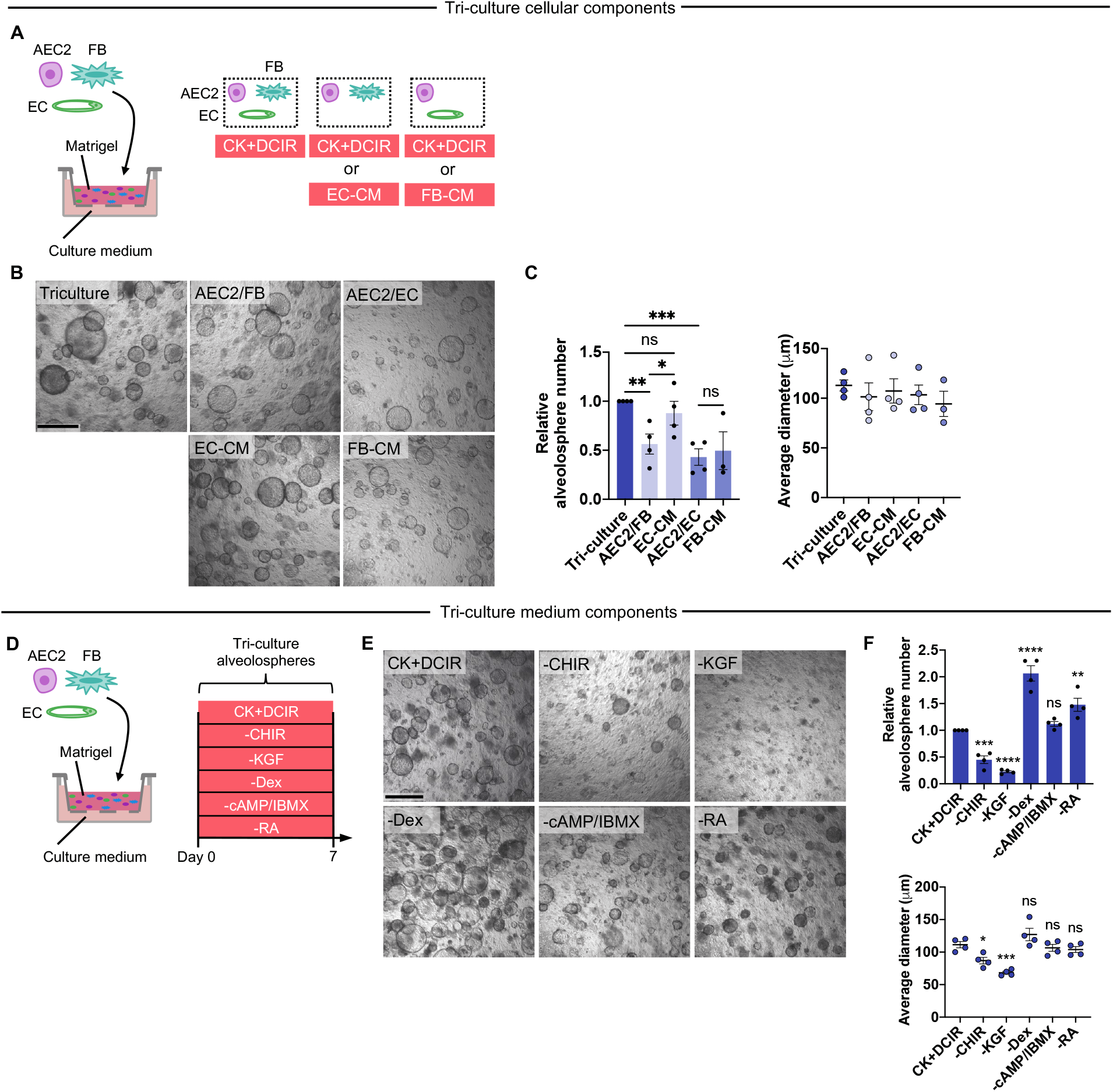
Contribution of tri-culture cellular and medium components to alveolosphere formation. (**A**) Design of conditioned medium (CM) alveolosphere assay. The same cell populations used for whole lung engineered cultures were suspended in Matrigel and cultured for 7 days with either CK+DCIR medium or conditioned medium (CM). (**B** and **C**) Phase contrast images (B) and quantification (C) of alveolospheres from assay in (A). *n* = 3: FB-CM; *n* = 4: all other groups. (C): Mixed-effects analysis with Holm-Sidak’s multiple comparisons tests. (**D**) Schematic of 3D alveolosphere assay to test the contribution of individual culture medium supplements to AEC2 growth in the tri-culture setting. The same cell populations used for whole lung engineered cultures were suspended in Matrigel and cultured for 7 days in complete CK+DCIR medium, or with CK+DCIR with the specified medium component(s) excluded. CHIR, CHIR99021. KGF, keratinocyte growth factor. Dex, dexamethasone. cAMP, cyclic AMP. IBMX, 3-isobutyl-1-methylxanthine. RA, retinoic acid. (**E** and **F**) Phase contrast images (E) and quantification (F) of alveolospheres from assay in (D). *n* = 4. (F): Repeated measures one-way ANOVA with Holm-Sidak’s multiple comparisons test. Error bars: mean ± SEM. *P* values given for comparison versus Tri-culture/CK+DCIR control. ns, not significant, \**P* < 0.05, \*\**P* < 0.01, \*\*\**P* < 0.001, \*\*\*\**P* < 0.0001. Scale bars, 500 μm.

**Extended Data Fig. 4.**
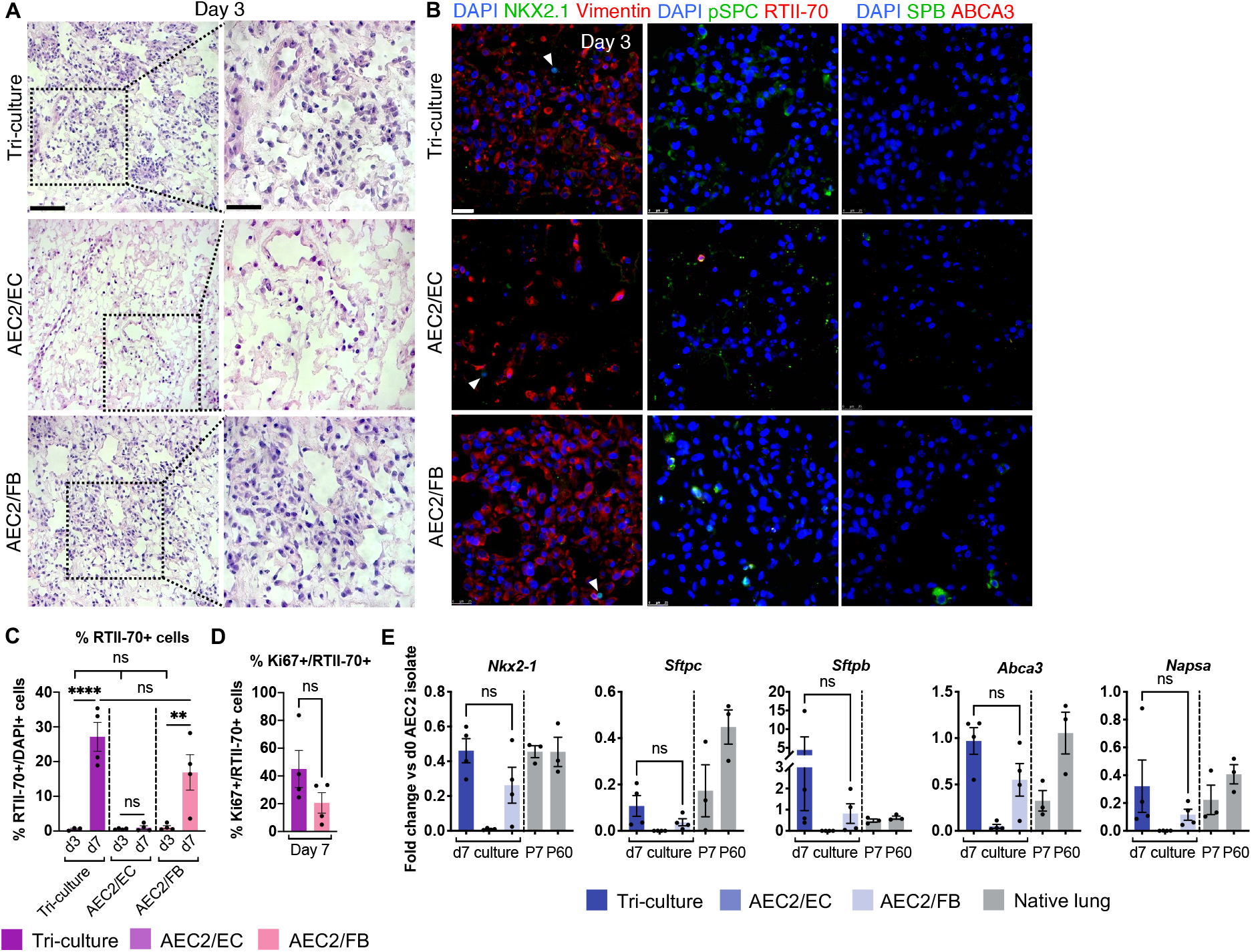
Fibroblasts are essential for AEC2 growth in decellularized lung scaffolds. (**A**) H&E of AEC2/FB, AEC2/EC, and tri-culture engineered lungs on day 3 after epithelial seeding. The relatively compressed tissue architecture of these samples reflects the difficulty of inflating just a portion of a lung lobe, without clear airway access. Scale bar, 100 μm. Magnified region, 50 μm. (**B**) Immunostaining for epithelial transcription factor NKX2.1 and AEC2 markers pSPC, RTII-70, SPB, and ABCA3 in day 3 engineered lungs, showing rare AEC2s at this timepoint across all conditions. Arrowheads, NKX2.1^+^ nuclei. Scale bar, 25 μm. (**C** and **D**) Quantification of percentage total (C) and proliferating (D) RTII-70+ cells in engineered lungs; *n* = 3 lungs: d3 tri-culture; *n* = 4: all other groups. (**E**) qRT-PCR of AEC2 gene expression in day 7 engineered lungs. Gene expression in native P7 and P60 rat lungs is normalized to the average expression in day 0 AEC2 isolates and shown for approximate comparison only. *n* = 4 lungs: engineered lungs; *n* = 3: P7 and P60. Error bars: mean ± SEM. (C): One-way ANOVA with Holm-Sidak’s multiple comparisons test. (D and E): Unpaired two-tailed *t*-test. ns, not significant, \*\**P* < 0.01, \*\*\*\**P* < 0.0001.

**Extended Data Fig. 5.**
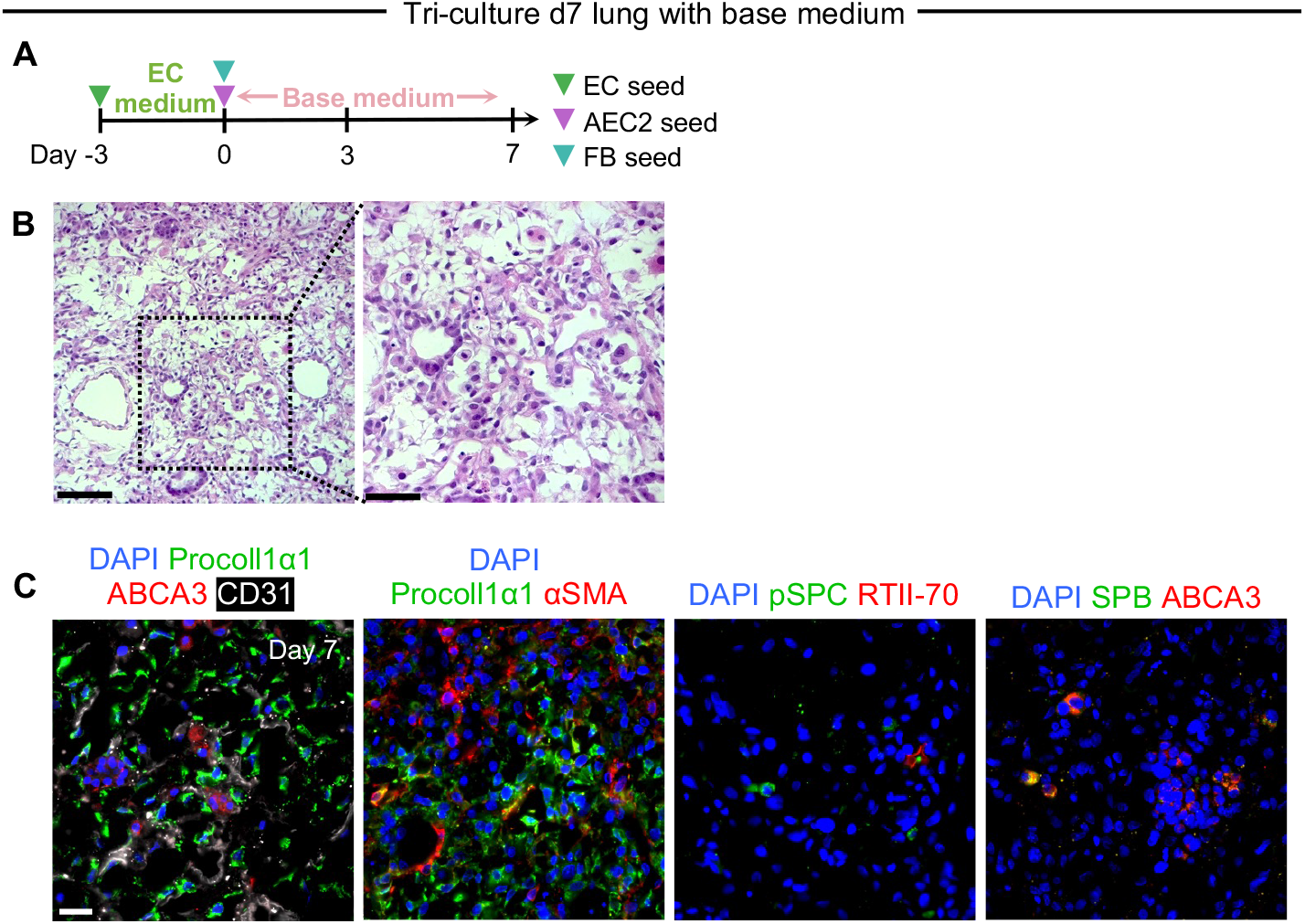
Endogenous tri-culture signaling in base medium is insufficient for organized scaffold repopulation with AEC2s. (**A**) Timeline for tri-culture engineered lung cultured without CK+DCIR medium additives (*i.e.* with CK+DCIR base medium only after day 0); *n* = 1. (**B**) H&E staining of tri-culture lung cultured with base medium only reveals disorganized repopulation and few cuboidal epithelial-like cells at 7 days. Scale bars: main image, 100 μm. Magnified region, 50 μm. (**C**) Immunostaining of tri-culture lung with base medium shows few scattered pSPC^+^, RTII-70^+^, or SPB^+^/ABCA3^+^ AEC2s within a disorganized stroma of CD31^+^ ECs and procoll1α1^+^ and/or αSMA^+^ FBs. Scale bars, 25 μm.

**Extended Data Fig. 6.**
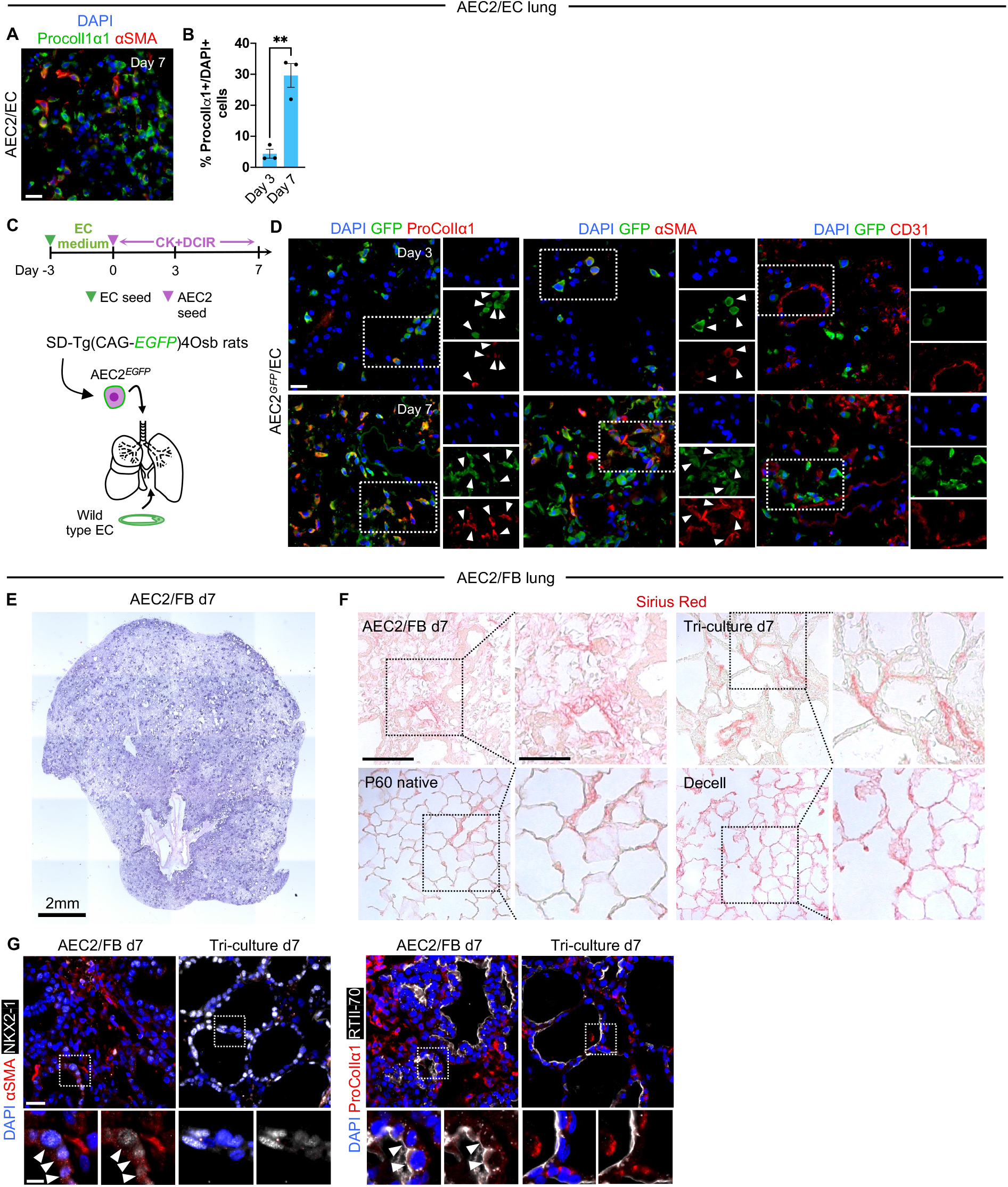
Additional characterization of AEC2/EC and AEC2/FB engineered lungs. (**A**) Immunofluorescent staining for procollagen Iα1 (ProColIα1) and alpha smooth muscle actin (αSMA) in day 7 AEC2/EC lungs. (**B**) Quantification of ProColIα1^+^ cell numbers in AEC2/EC engineered lungs, showing an increase in the number of FB-like cells over time (*n* = 3). Error bars: mean ± SEM. Unpaired two-tailed *t*-test: \*\**P* < 0.01. (**C**) Timeline for AEC2*^EGFP^*/EC lung culture, in which AEC2s isolated from *EGFP*^+^ rats were cultured together with wild-type ECs (*n* = 1). Under these conditions, all GFP^+^ cells in the resulting lung culture will have arisen from the AEC2 isolate. (**D**) Immunofluorescent staining of day 3 and 7 AEC2*^EGFP^* /EC lungs showing GFP^+^ staining co-localizing with FB markers (ProColIα1 and αSMA – see arrowheads) but not with CD31. All scale bars, 25 μm. (**E**) Whole lobe H&E staining of d7 AEC2/FB engineered lung. (**F**) Representative images of Sirius Red staining for collagen in engineered, native, and decellularized lung. Note the disorganized collagen deposition and/or remodeling, with accompanying disruption of the alveolar meshwork, in AEC/FB engineered lung. Scale bar, 100 μm. Magnified region, 50 μm. (**G**) Immunostaining demonstrating αSMA and ProColIα1 protein expression in d7 AEC2/FB, but not tri-culture, engineered lung epithelium. Arrowheads, epithelial cells positive for αSMA or proColIα1. Scale bars: main image, 25 μm; magnified region, 10 μm.

**Extended Data Fig. 7.**
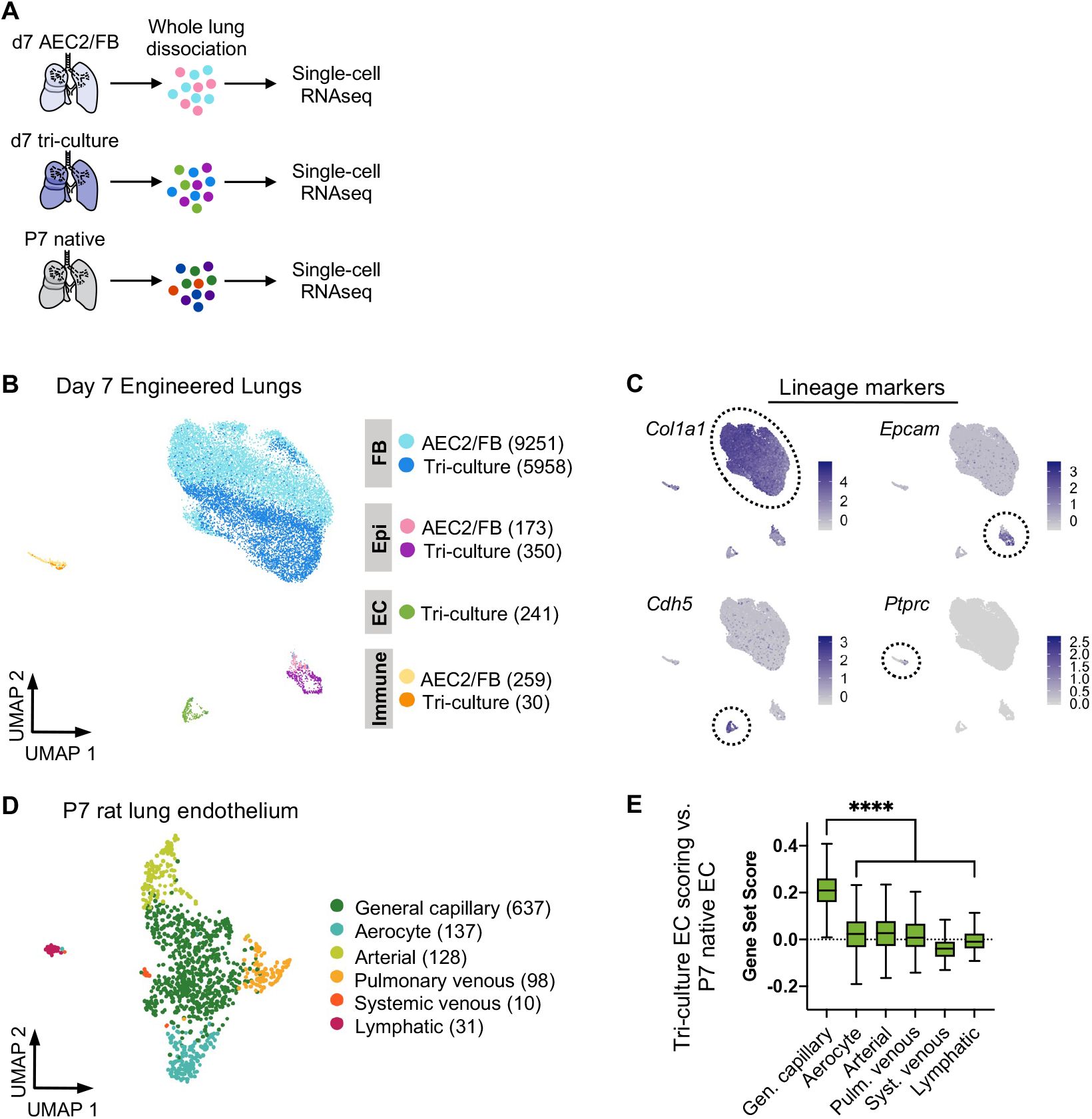
Overview of engineered lung scRNAseq data. (**A**) Schematic of scRNAseq analysis of engineered and native lungs. (**B**) UMAP embedding of scRNAseq data for AEC2/FB and tri-culture day 7 engineered lungs. Samples were clustered independently but are integrated here for aid of visualization. Clusters are colored by lung culture and cell lineage. (**C**) Expression data for example markers used to assign cells to fibroblast (FB), epithelial (epi), endothelial cell (EC), or immune lineages in engineered lungs, projected onto scRNAseq data from (B). Dotted ovals encircle clusters positive for the indicated marker. Note that the scRNAseq data revealed a cluster of *Ptprc*^+^ immune cells in each lung, most likely derived from the AEC2 cell isolate (see Extended Data Fig. 1B,C). While we cannot exclude a role for these immune cells in the observed lung phenotypes, they were not included in the analysis in this study. (**D**) UMAP embedding of scRNAseq data for P7 rat lung endothelial cells, extracted from the full P7 data in fig. S1E and re-clustered. (**E**) Scoring of individual tri-culture endothelial cells for expression of top markers from each of the P7 endothelial populations shown in (D). Scores > 0 indicate enriched expression compared to random gene sets. Boxplots: center indicates median, box limits indicate 1^st^ and 3^rd^ quartiles, whiskers indicate min and max (outliers not shown). Kruskal-Wallis test with Dunn’s post-test. General capillary score vs each other individual score, \*\*\*\**P* < 0.0001. (B and D) Numbers in parentheses indicate cell numbers per cluster after filtering.

**Extended Data Fig. 8.**
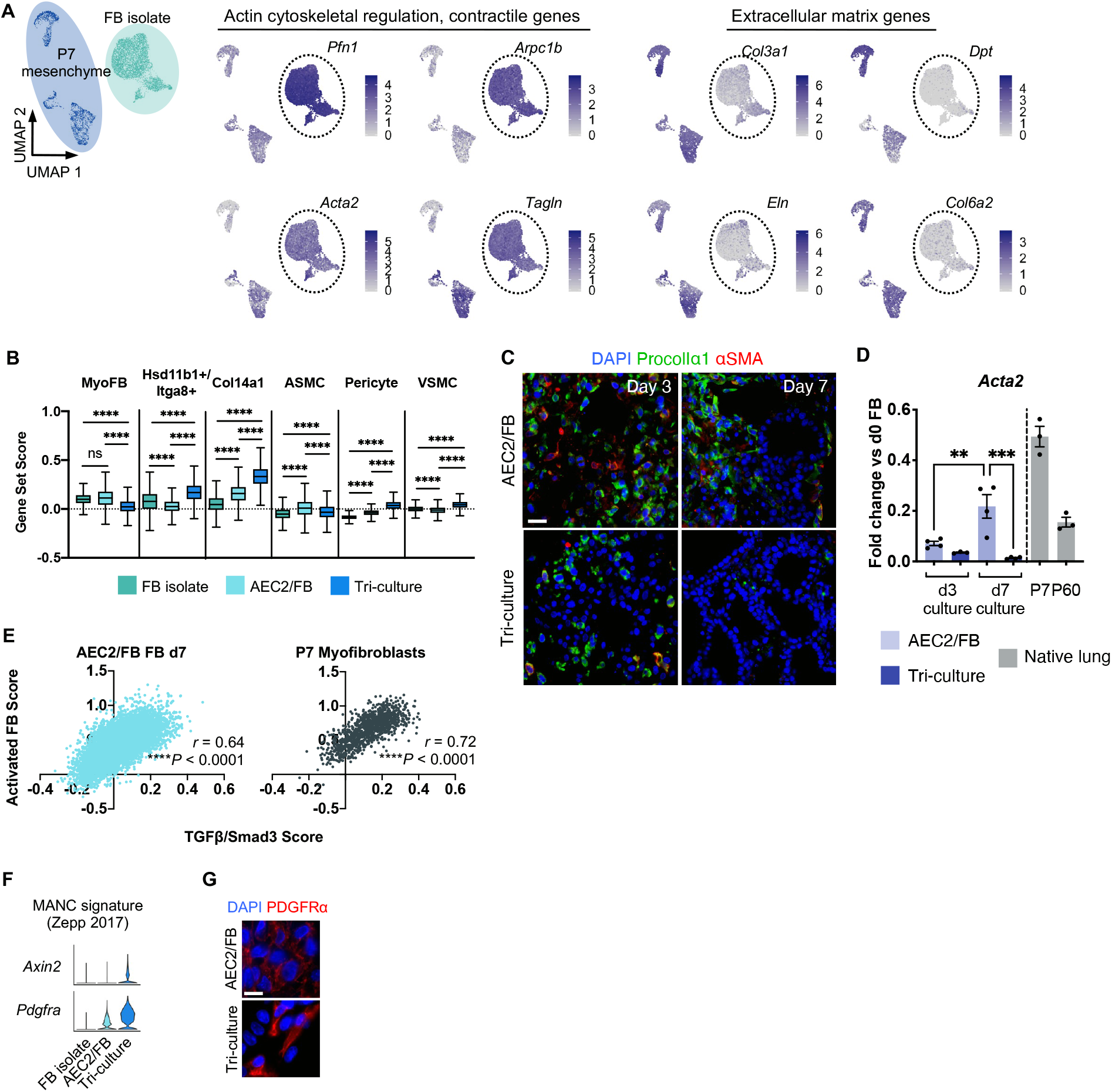
Additional characterization of the FB isolate and engineered lung FB populations. (**A**) Expression of select contractile (middle) and extracellular matrix genes (right), projected onto scRNAseq data of P7 mesenchyme (see Extended Data Fig. 2F) and the FB isolate mesenchyme (see Extended Data Fig. 2G), demonstrating an overall shift toward a contractile and pro-migratory phenotype in the cultured FB isolate. UMAP with cell identities is shown in left of panel; populations have been merged for purposes of expression level comparison. Dotted ovals encircle clusters positive for the indicated marker. (**B**) Scoring of individual FBs for expression of P7 native mesenchymal cluster gene sets. MyoFB, myofibroblast. ASMC, airway smooth muscle cell. VSMC, vascular smooth muscle cell. Boxplots: center indicates median, box limits indicate 1^st^ and 3^rd^ quartiles, whiskers indicate min and max (outliers not shown). Kruskal-Wallis test with Dunn’s post-test. ns, not significant; \*\*\*\**P* < 0.0001. (**C**) Immunostaining of day 3 and day 7 engineered lungs for procollagen Iα1 (ProcolIα1) and αSMA (gene *Acta2*). Scale bar, 25 μm. (**D**) qRT-PCR for *Acta2* in engineered lung tissue. *n* = 3, d3 tri-culture, P7 and P60 native; *n* = 4, all other groups. Native gene expression is normalized to the average expression in day 0 FB isolates and shown for approximate comparison only. Error bars indicate the mean ± SEM. One-way ANOVA with Holm-Sidak’s multiple comparisons test. \*\**P* < 0.01, \*\*\**P* < 0.001. (**E**) Scatterplots with associated Spearman correlation coefficients *r* showing moderate correlations between TGFβ/Smad3 target gene expression and activation scores in AEC2/FB FBs and P7 myofibroblasts. (**F**) Violin plots of expression of mesenchymal alveolar niche cell (MANC) signature genes *Axin2* and *Pdgfra* in the cultured FB isolate and engineered lung FBs. (**G**) Immunostaining of PDGFRα protein expression in d7 engineered lung FBs. Scale bar, 10 μm.

**Extended Data Fig. 9.**
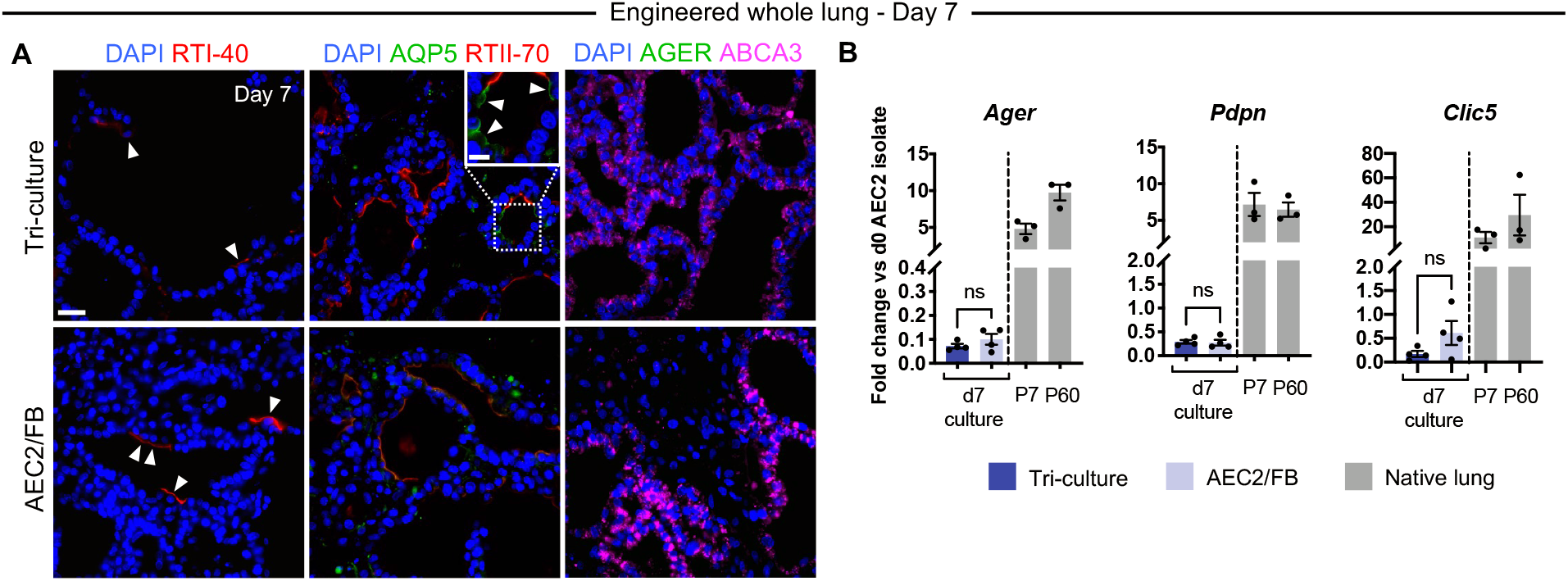
AEC2/FB and tri-culture lungs cultured with CK+DCIR medium show minimal evidence of AEC2 differentiation to AEC1s. (**A**) Immunostaining for AEC1 markers RTI-40, AQP5, and AGER in d7 engineered lungs. Arrowheads, cells staining positive for AEC1 markers. Scale bars: main image, 25 μm; magnified region, 10 μm. (**B**) qRT-PCR of AEC1 gene expression in d7 engineered lungs; *n* = 4 lungs. Gene expression in native P7 and P60 rat lungs is normalized to the average expression in day 0 AEC2 isolates and shown for approximate comparison only; *n* = 3 lungs. Error bars: mean ± SEM. Unpaired two-tailed *t*-test. ns, not significant.

**Extended Data Fig. 10.**
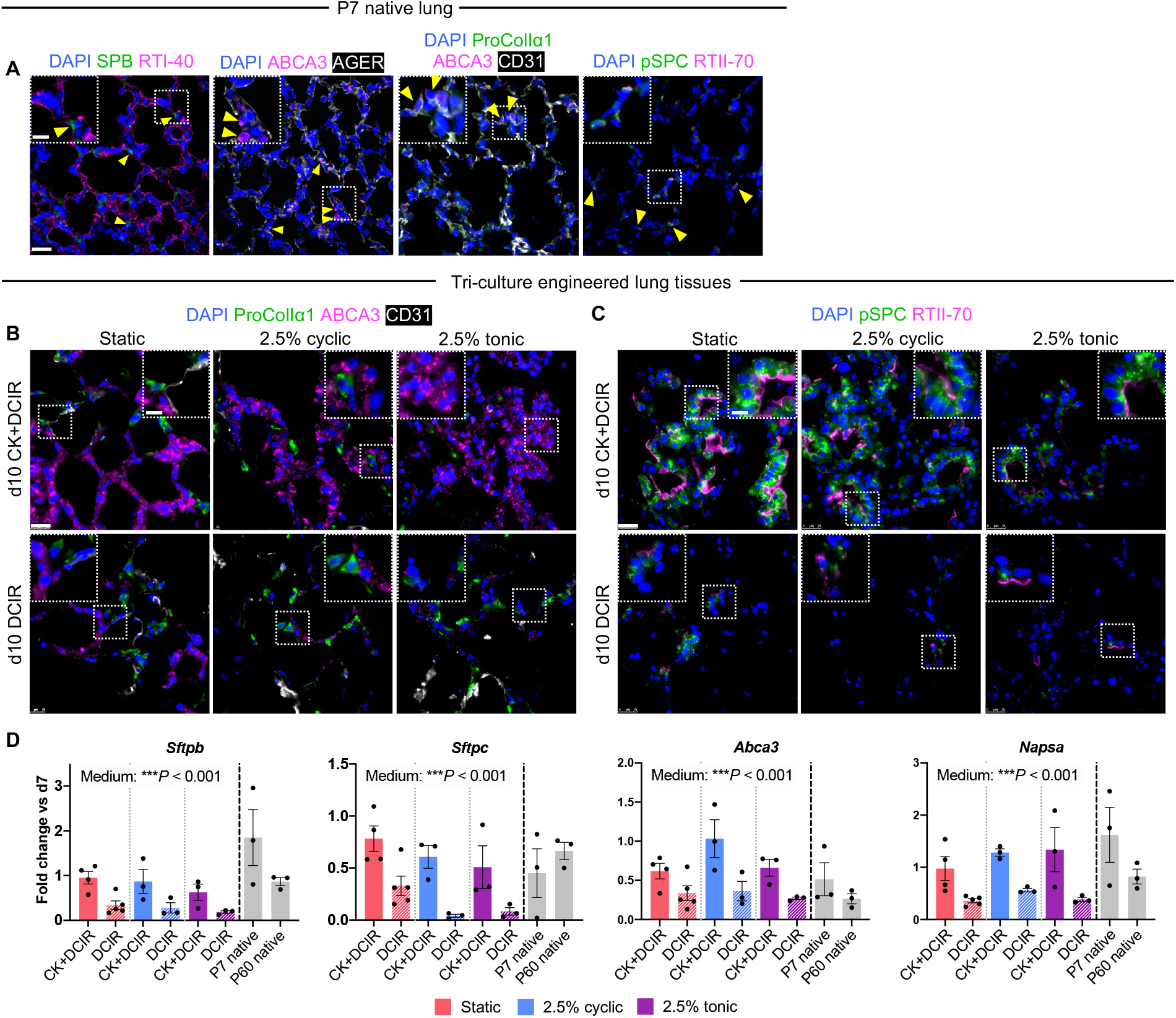
CHIR and KGF withdrawal is associated with a loss of AEC2 phenotype in tri-culture ELTs. (**A**) Immunostaining of P7 native lung as controls for ELTs in Fig. 6, C and D and this figure (B and C). Note that imaging parameters used for the native tissues differ from those used for the ELTs, as higher exposures were generally required to visualize the protein staining patterns in native lung. Arrowheads, AEC2s localized to alveolar corners. (**B** and **C**) Immunostaining for AEC2s (ABCA3, pSPC, RTII-70), FBs (ProColIα1) and ECs (CD31) in tri-culture d10 ELTs. (**D**) qRT-PCR of AEC2 gene expression in d10 ELTs, normalized to their respective d7 control tissues. Native gene expression is normalized to the average expression in d7 tissues and shown for approximate comparison only. Static/CK+DCIR: *n* = 4; static/DCIR: *n* = 5; all other conditions: *n* = 3. Error bars: mean ± SEM. Text indicates significance by two-way ANOVA for medium main effect (“medium”). Scale bars: main image, 25 μm; magnified region, 10 μm.

