## Supplementary material for "*In vitro* engineering of the lung alveolus": Summary of supplemental tables, Supplementary Tables 4 and 5

**Supplementary information**

List of Supplementary Tables:

**Supplementary Table 1: Engineered lung scRNAseq cluster markers (separate Excel spreadsheet)**
Single-cell RNAseq cluster markers for day 7 AEC2/FB engineered lung (tab 1) and day 7 tri-culture engineered lung (tab 2).

**Supplementary Table 2: Engineered lung connectome data (separate Excel spreadsheet)**
Ligand-receptor connectomes for day 7 AEC2/FB engineered lung (tab 1) and day 7 tri-culture engineered lung (tab 2).

**Supplementary Table 3: scRNAseq fibroblast analysis (separate Excel spreadsheet)**
Single-cell RNAseq cluster markers for the cultured FB isolate, engineered lung FBs, and P7 native FBs (tab 1); differentially expressed genes (DEGs, tab 2) with associated Hallmark pathway enrichment (tab 3) enriched in the FB isolate and day 7 AEC2/FB FBs, as compared to day 7 tri-culture FBs (corresponds to Fig. 4C); and biological process (BP) enrichment in day 7 AEC2/FB FBs (tab 4) and day 7 tri-culture FBs (tab 5) (corresponds to Fig. 4D).

**Supplementary Table 4: Antibodies and dilutions used for immunofluorescent staining.**

**Supplementary Table 5: qRT-PCR primer sequences.**

**Supplementary Table 6: Gene sets used for cell scoring (separate Excel spreadsheet)**

**Supplementary Table 4: Antibodies and dilutions used for immunofluorescent staining**

| Primary Antibody | Host species | Supplier | Product number | Dilution | Clone |
|---|---|---|---|---|---|
| Anti-αSMA | Mouse | Dako | M0851 | 1:2000 | 1A4 |
| Anti-ABCA3 | Mouse | Abcam | ab24751 | 1:50 | 3C9 |
| Anti-AGER | Goat | R&D Systems | AF1145 | 1:200 | N/A |
| Anti-AQP5 | Rabbit | Millipore | AB3559 | 1:500 | N/A |
| Anti-CD31 | Goat | R&D Systems | AF3628 | 5 μg/mL | N/A |
| Anti-GFP | Mouse | Santa Cruz | sc-9996 | 1:50 | B-2 |
| Anti-GFP | Rabbit | Abcam | ab290 | 1:5000 | N/A |
| Anti-Laminin | Rabbit | Abcam | ab11575 | 1:200 | N/A |
| Anti-NKX2.1 | Rabbit | Abcam | ab76013 | 1:100 | EP1584Y |
| Anti-PDGFRα | Rabbit | Abcam | ab203491 | 1:250 | EPR22059-270 |
| Anti-procollagenIα1 | Rabbit | Rockland | 600-401-D19 | 1:50 | N/A |
| Anti-pSPC | Rabbit | Millipore | AB3786 | 1:1000 | N/A |
| Anti-RTI-40 | Mouse | Terrace Biotech | TB-11ART1-40 | 1:200 | N/A |
| Anti-RTII-70 | Mouse | Terrace Biotech | TB-44ART2-70 | 1:40 | N/A |
| Anti-Smad3-pS423/pS425 | Rabbit | Rockland | 600-401-919 | 1:400 | N/A |
| Anti-SPB | Rabbit | Santa Cruz | sc-13978 | 1:50 | H-300 |
| Anti-TGFβ1 | Rabbit | Abcam | ab92486 | 1:200 | N/A |
| Anti-Vimentin | Mouse | Abcam | ab8069 | 1:200 | V9 |
| Secondary Antibody | | | | | |
| Anti-goat IgG, Alexa Fluor 488 | Chicken | Invitrogen | A21467 | 1:500 | N/A |
| Anti-goat IgG, Alexa Fluor 555 | Donkey | Invitrogen | A21432 | 1:500 | N/A |
| Anti-goat IgG, Alexa Fluor 647 | Donkey | Invitrogen | A-21447 | 1:500 | N/A |
| Anti-mouse IgG, Alexa Fluor 555 | Goat | Invitrogen | A21424 | 1:500 | N/A |
| Anti-mouse IgG, Alexa Fluor 568 | Donkey | Invitrogen | A10037 | 1:500 | N/A |
| Anti-mouse IgG, Alexa Fluor 647 | Goat | Invitrogen | A-21235 | 1:500 | N/A |
| Anti-rabbit IgG, Alexa Fluor 488 | Chicken | Invitrogen | A21441 | 1:500 | N/A |

| | | | | | |
|---|---|---|---|---|---|
| Anti-rabbit IgG, Alexa Fluor 488 | Goat | Invitrogen | A11034 | 1:500 | N/A |
| Anti-rabbit IgG, Alexa Fluor 555 | Goat | Invitrogen | A21429 | 1:500 | N/A |
| Anti-rabbit IgG Alexa Fluor 647 | Donkey | Invitrogen | A31573 | 1:500 | N/A |

**Supplementary Table 5: qRT-PCR primer sequences.**

| Rat Primer | Forward | Reverse |
|---|---|---|
| *Abca3* | GAGGTCTTCCTTCGGGTTGG | GTCCATCACCCCACACAAGT |
| *Acta2* | GCTTTGCTGGTGATGATGCT | GATCCCTCTCTTGCTCTGC |
| *Actb* | GCAGGAGTACGATGAGTCCG | ACGCAGCTCAGTAACAGTCC |
| *Ager* | AGAAACCGGTGATGAAGGACA | GGTTGTCGTTTTCGCCACAG |
| *B2m* | CCGTGATCTTTCTGGTGCTT | ATTTGAGGTGGGTGGAACTG |
| *Clic5* | CTGGCCGACTGCAATCTACT | GTGAACTCGTCCCGTGCATA |
| *Napsa* | CAGGTCCACATGCAGAGTGT | GCCTTATTCAAGGCCCGGAT |
| *Nxk2-1* | TGCTTTATGGTCGGACCTGG | TTGCGGAGGGTAGAGGGAAA |
| *Pdpn* | AGTGTTGCTCTGGGTTTTGG | GGGTTCACCATGTCATCTCC |
| *Sftpb* | CCTGGCTGAGCGTTACACA | TTCAATCAGAGGCTCCAGAG |
| *Sftpc* | CTCCTGACCGCCTATAAGC | TGGCCTGGAAGTTCTTGAAT |
